# Aging-Dependent Loss of Connectivity in Alzheimer’s Model Mice with Rescue by mGluR5 Modulator

**DOI:** 10.1101/2023.12.15.571715

**Authors:** Francesca Mandino, Xilin Shen, Gabriel Desrosiers-Gregoire, David O’Connor, Bandhan Mukherjee, Ashley Owens, An Qu, John Onofrey, Xenophon Papademetris, M. Mallar Chakravarty, Stephen M Strittmatter, Evelyn MR Lake

**Author notes:** Co-senior authorship (Stephen M Strittmatter, Evelyn MR Lake). **Correspondence**: Stephen M. Strittmatter & Evelyn M.R. Lake Anlyan Center, 300 Cedar St, New Haven, CT 06519 or.

## Abstract

Amyloid accumulation in Alzheimer’s disease (AD) is associated with synaptic damage and altered connectivity in brain networks. While measures of amyloid accumulation and biochemical changes in mouse models have utility for translational studies of certain therapeutics, preclinical analysis of altered brain connectivity using clinically relevant fMRI measures has not been well developed for agents intended to improve neural networks. Here, we conduct a longitudinal study in a double knock-in mouse model for AD (*App^NL-G-F^/hMapt*), monitoring brain connectivity by means of resting-state fMRI. While the 4-month-old AD mice are indistinguishable from wild-type controls (WT), decreased connectivity in the default-mode network is significant for the AD mice relative to WT mice by 6 months of age and is pronounced by 9 months of age. In a second cohort of 20-month-old mice with persistent functional connectivity deficits for AD relative to WT, we assess the impact of two-months of oral treatment with a silent allosteric modulator of mGluR5 (BMS-984923) known to rescue synaptic density. Functional connectivity deficits in the aged AD mice are reversed by the mGluR5-directed treatment. The longitudinal application of fMRI has enabled us to define the preclinical time trajectory of AD-related changes in functional connectivity, and to demonstrate a translatable metric for monitoring disease emergence, progression, and response to synapse-rescuing treatment.

## Introduction

It can take decades for the hallmarks of Alzheimer’s disease (AD) to clinically manifest. Optimistically, this is evidence of a long therapeutic window during which treatment could interrupt or reverse disease processes, provided we have access to ways of identifying patients, staging disease, and effective therapeutics. Functional magnetic resonance imaging (fMRI) is a strong candidate for helping to address these needs, given that it is non-invasive and can assay the entire brain. BOLD (blood-oxygen-level dependent) fMRI yields measures of functional connectivity (FC, or inter-regional BOLD signal synchrony). In AD, robust changes in FC are observed, with brain-network specificity, years prior to the clinical manifestation of neurodegenerative signatures of disease.^1–8^ Changes in FC progress in concert with AD-related declines in executive function and cognition, lending confidence to the notion that aberrant FC is a harbinger of neurodegeneration. Further, changes in FC follow a similar spatial trajectory as subsequent brain atrophy, Aβ-pathology, neurofibrillary tangle deposition,^9,10^ and synapse losses.^11–14^ In sum, BOLD-fMRI FC may be sensitive to the varied stages of AD emergence and progression. However, despite tremendous effort, and investment, human fMRI research has yet to produce actionable neuroimaging markers of AD.^4^ In part, this is due to the long time-scale and complexity of AD pathology, the difficulty of relating fMRI markers to underlying mechanisms, and substantial heterogeneity within the population.

To help address these gaps, the application of BOLD-fMRI in animal models of AD is fundamental for improving our understanding of the imaging correlates of disease and treatment response.^15–17^ Studies in animals provide a well-controlled environment for measuring disease phenotypes and testing therapeutics on a tractable timescale, as well as the freedom to obtain invasive measurements. The short lifespan, low-cost, and genetic malleability of mice, together with recent improvements in rodent BOLD-fMRI data quality and processing techniques^18–22^ (including inter-species translation),^23,24^ mean the field is well-positioned to make new inroads into AD research. Advances in how we model AD in mice, where it has been hard to faithfully recapitulate human disease,^25^ lends further cause for enthusiasm. Here, we use the double-knock-in (DKI) *App^NL-G-F^/hMapt* (amyloid-precursor-protein/human microtubule-associated protein tau) mouse model,^26,27^ which critically avoids overexpression artifacts. With this model, we seek to establish BOLD-fMRI FC correlates of AD emergence, progression, and treatment response. We collect longitudinal resting-state BOLD-fMRI data spanning a wide-range of AD-relevant timepoints, from 4 to 22 months of age (M) or the rough equivalent of 20-70 years of age in humans.^28^ In late-stage disease (20-22M), we treat a subgroup of mice with a metabotropic glutamate receptor 5 (mGluR5) silent allosteric modulator (BMS-984923).^29^ Vitally, treatment is given during very late-stage disease which is in stark contrast to existing anti-amyloid drugs which have only shown efficacy during early (asymptomatic) illness.^30^ Further, this compound is known to modify AD course by rescuing synapses and improving memory,^31^ and has recently entered clinical trials (ClinicalTrials.gov; NCT05804383). However, the treatment-elicited effects on BOLD-fMRI FC are currently unknown. We seek to gain this knowledge given the potential usefulness of BOLD-fMRI FC for monitoring disease and treatment response.

We find that FC in AD mice follows a distinct age-related trajectory relative to wild-type (WT) littermates. In AD mice, changes emerge later and show more widespread decreases in FC with time, compared to WT mice. The age and AD-related brain networks we identify using data-driven approaches show significant enrichment for established circuits including the default mode and lateral cortical networks (DMN and LCN, respectively).^32,33^ Further, oral treatment with BMS-984923 rescues FC deficits in AD mice. In sum, this work identifies age- and AD-related phenotypes in BOLD-fMRI data across the mouse lifespan in a relevant model of disease.^26,27^ We also find timely evidence that BOLD-fMRI FC may prove useful for assessing the effectiveness of a novel synapse-rescuing treatment.

## Materials and methods

All procedures are approved by the Yale Institutional Animal Care and Use Committee (IACUC) and follow the National Institute of Health Guide for the Care and Use of Laboratory Animals. Both biological sexes are included. Mice are group housed on a 12-hour light/dark cycle with *ad libitum* food and water. Cages are individually ventilated as well as temperature and humidity controlled.

### Model of Alzheimer’s disease

Among available AD models there are amyloid-precursor-protein (App) knock-in (KI) strains.^26,27^ Here, we use *App^NL-G-F^* KI mice where aβ-containing exons from human APP with Swedish, Arctic, and Iberian mutations are included^26^. In addition, mice are crossed with a strain in which the human microtubule-associated protein tau (*MAPT*) locus, with its full sequence and splice sites, replaces the murine locus to generate *App^NL-G-F^/hMapt* homozygous double-knock-in animals.^26^ The strain has been back-crossed with C57Bl6J for >10 generations, with strain matching confirmed by genetic markers. Male and female AD and wild-type (WT) littermate controls are included.

### Dataset overview

A total of N=74 animals are included from across three experiments (**Table 1**). In the first experiment, N=27 animals undergo transverse sinus injections on postnatal day zero (P_0_) to induce whole-brain fluorescence. These mice then undergo head-post surgery at 3M and multimodal imaging at 4, 6, and 9M where wide-field calcium (WF-Ca^2+^) and fMRI data are collected simultaneously.^34,35^ The WF-Ca^2+^ imaging data are the subject matter of a separate publication. In the second experiment, N=16 animals undergo head-post surgery at 5 or 8M and unimodal imaging (fMRI-only) at 6 or 9M, respectively. These data are collected to address attrition in the first experiment. In the final experiment, N=30 animals undergo head-post surgery at 19M and unimodal imaging (fMRI-only) at 20 and 22M. Between these imaging sessions, AD mice are split between a treatment (N=11) and vehicle (N=9) subgroup.

**Table 1.**
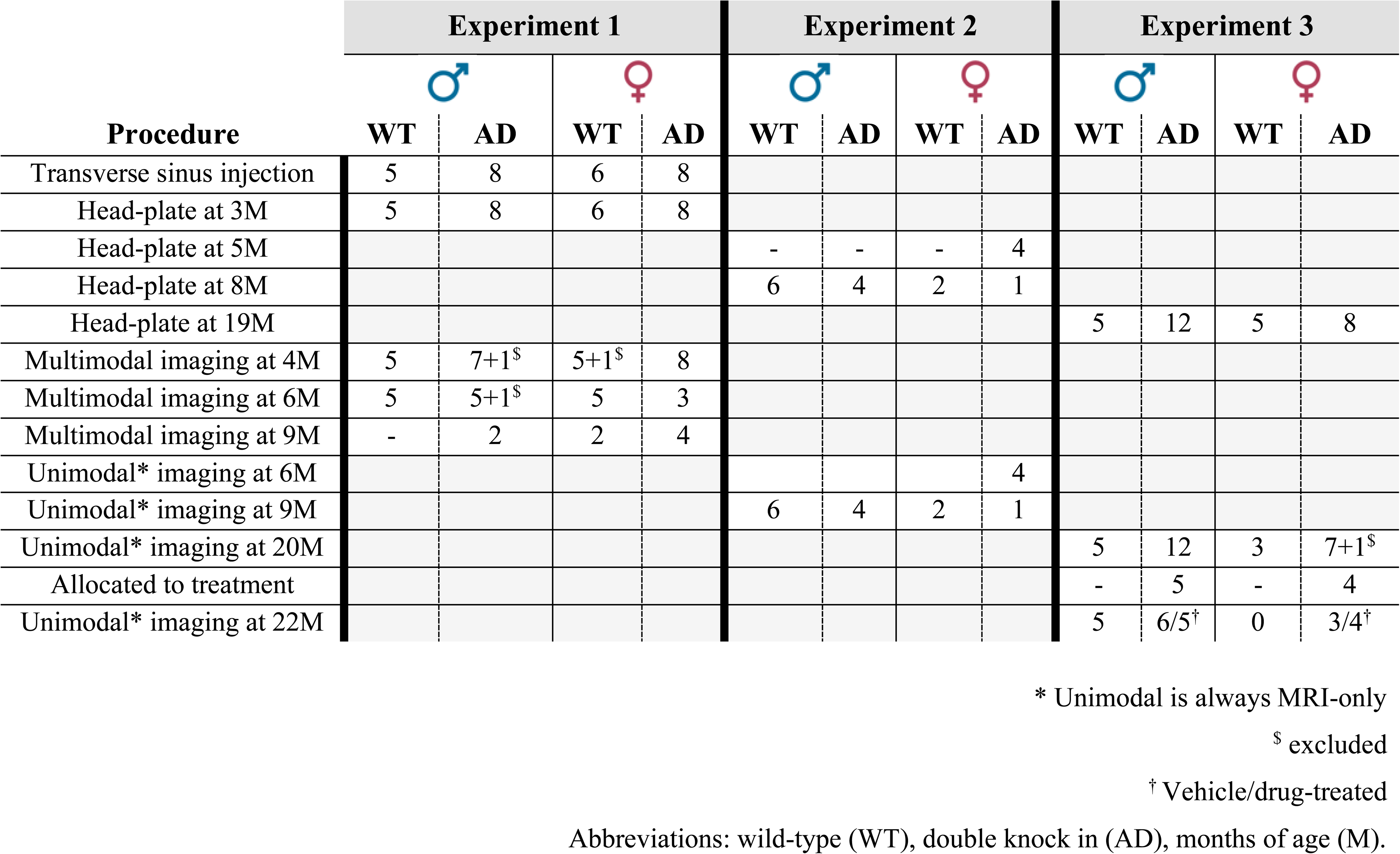
Overview of datasets.

### Head-implant surgery

Surgical details are provided in the **Supplementary Material**.

### Data acquisition

All mice undergo imaging under anaesthesia (free-breathing low-dose isoflurane, 0.5-0.75%, in 50/50 medical air/O_2_). The MRI data are acquired on an 11.7T preclinical magnet (Bruker, Billerica, MA), using ParaVision version 6.0.1 software. Body temperature is monitored (Neoptix fiber), maintained with a circulating water bath at 36.6-37°C, and recorded (Spike2, Cambridge Electronic Design Limited, Master-8 A.M.P.I.). Breathing rate is measured using a foam respiration pad (SAM-32, Starr).

#### Structural MRI data acquisition

At every imaging session, three structural images are acquired, only one is used for data registration purposes. (1) A whole-brain isotropic 3D-image using a MSME sequence, with 0.2×0.2×0.2mm^3^ resolution, TR/TE of 5500/20ms, 78 slices, 2 averages (11mins and 44s). For the remainder two structural images see **Supplementary Methods**.

#### Functional MRI data acquisition

Data are acquired using a gradient-echo, echo-planar-imaging (GE-EPI) sequence with a TR/TE of 1.8s/11ms. Data are collected across 25 slices without gaps at 0.31×0.31×0.31mm^3^; resulting in near whole-brain coverage. Each run has 334 repetitions, corresponding to 10min of data per run. We acquire 3 runs/session for a total of 30min of data per mouse per imaging session.

### Data pre-processing and analysis

#### Functional MRI pre-processing and registration to common space

Data registration and pre-processing is performed using RABIES (Rodent automated BOLD improvement of EPI sequences) v0.4.8 (https://rabies.readthedocs.io/en/stable/).^36^ All functional and isotropic MSME structural scans (from all imaging sessions and experiments, **Table 1**) are input and pre-processed together. Slice time correction is first applied on native space timeseries, and head motion parameters are estimated.

The isotropic MSME structural images (one per mouse per session) are corrected for inhomogeneity (N3 nonparametric nonuniform intensity normalization)^37,38^, non-linearly registered and then averaged to create a within-sample template. The within-sample template is non-linearly registered to an out-of-sample template we created from N=162 datasets (collected on the same scanner from N=64 mice from a separate longitudinal study which uses the same isotropic MSME sequence), see **Supplementary Material**. The out-of-sample template is registered to the Allen Atlas reference space CCfv3^39,40^. For more details on pre-processing steps, registrations and denoising see **Supplementary Material**.

#### Exclusion criteria

Of all N=74, N=6 animals and n=2 runs are completely excluded. N=1 for enlarged ventricles, N=1 for a brain tumour, N=2 for anaesthesia malfunction (i.e., tubes were improperly connected, N=1, 4M and N=1, 6M), and N=2 due to implant detachment (caused by a glass coating that was not present in other surgeries, see **Supplementary Material**). An additional n=2 runs were excluded (n=1, 6M and n=1, 9M) due to image registration failures. More details are given in the **Supplementary Material**. Further, data quality was inspected using the RABIES *--data diagnosis* toolkit.^36^ Finally, no data are excluded due to a low signal to noise ratio or motion (FD>0.075mm in over 1/3^rd^ of frames per run).

#### Connectomes

The 172 regions of interest (ROIs) from the Allen Atlas and MRI template space (**Fig. S1**) are shared as part of BioImage Suite, https://bioimagesuiteweb.github.io/alphaapp/connviewer.html?species=mouse#. The ROIs are selected based on the resolution of our fMRI data, two regions are excluded for visualization purposes for the matrices. To compute connectomes, BOLD signals within each ROI are averaged, and the inter-regional Pearson’s correlation computed. Results are computed for each run using data in the out-of-sample template space. For a description of the data-derived networks and the triple network model see **Supplementary Material**.

#### Treatment

Drug synthesis, incorporation into diet, feeding, and validation are based on previous work. See summary in **Supplementary Material**.

#### Statistics

Functional connectomes are computed using the Pearson’s correlation and converted to z-scores using the fisher r to z transform. Descriptive statistics are given as mean differences and 25/75^th^ percentiles unless stated otherwise, and graphically represented as box plots (MATLAB, *boxplot*). Statistical significance tests are computed in MATLAB (R2021bV5). Differences between individual edges are tested using a two-sample t-test (MATLAB, *ttest2*), and multiple comparison correction are applied using the Benjamini-Hochberg method. Furthermore, the significance of the derived networks are confirmed with permutation testing (×1,000 iterations) where the test statistics of the null distribution is the size of the network of a given comparison, and the group labels of the data are randomly shuffled.. The two sample t-test statistical threshold is set to p<0.05, two-tailed. The false discovery rate for the Benjamini-Hochberg test is set to be 0.05. The test shows that the true group labels lead to significantly more different edges than random labelling. For the 22M timepoint, a one-way ANOVA (MATLAB, *anova1*) is used to test if there is any pair of groups (WT_vehicle_ vs. AD_vehicle_, WT_vehicle_ vs AD_treated_, and AD_vehicle_ vs AD_treated_) that are significant different from each. A follow-up t-test is applied to identify which two groups are significantly different, results reported as above. The overlap of each couple of inputs to each data-derived network (**Fig. 2, 4, 5**) are assessed with Jaccard similarity test and compared to a null distribution (×1,000 iterations) to confirm the meaningfulness of the significant overlap found. Finally, the over/under enrichment (i.e., representation that is above chance) of the triple-network model with the data-derived networks is calculated with Fisher’s exact test as well as permutation testing (as described above, ×1,000 iterations).

### Data Availability

Upon reasonable request to the corresponding authors.

## Results

Male and female mice (N=74) from three longitudinal BOLD-fMRI experiments are included (**Table 1**). Prior to imaging, all animals undergo a minimally invasive surgical procedure,^34,35,41^ where a head-plate is affixed to the skull a month prior to any imaging (timeline, **Fig. 1**). Mice are imaged longitudinally at 4, 6 and 9M (experiment 1), at 6 and/or 9M (experiment 2), and at 20 and 22M (experiment 3). At all imaging timepoints, MRI data are collected from a double knock-in (DKI) mouse model of AD (*App^NL-G-F^/hMapt*)^26,27^ and WT littermate controls. After the 20M imaging timepoint, a subgroup of AD mice is randomly assigned to oral treatment with BMS-984923^29^ (AD_treated_) or regular chow (AD_vehicle_). Thirty-minutes of resting-state BOLD-fMRI data, together with structural images for data registration, are collected at each timepoint under low-dose isoflurane (0.5-0.75%). Data are processed and registered to a common space.^34–36^ A small fraction of the fMRI data (9%) are excluded based on quality control standards.^18,36,42,43^ Z-scored functional connectomes (brain-wide FC measurements) are computed using 172-regions from the Allen Atlas CCfv3^39,40^ (key, **Fig. S1**) and Pearson’s correlation. Average connectomes for each group at each timepoint are shown in **Fig. 1**. Stereotypical patterns including high FC between bilateral brain regions are observed. Differences between connectomes are displayed with and without thresholding for significance (*p<*0.05) at the edge-level and correcting for multiple comparisons (**Methods**). Permutation testing (by randomly shuffling datasets) is applied (×1,000 iterations). In all results, the test statistic of the null distribution is the number of edges (region-to-region measures of FC) remaining after correcting for multiple comparisons. Identified brain circuits that show significant effects are summarized as connectomes, vertical histograms, circle plots, and node-degree maps.

**Fig. 1.**
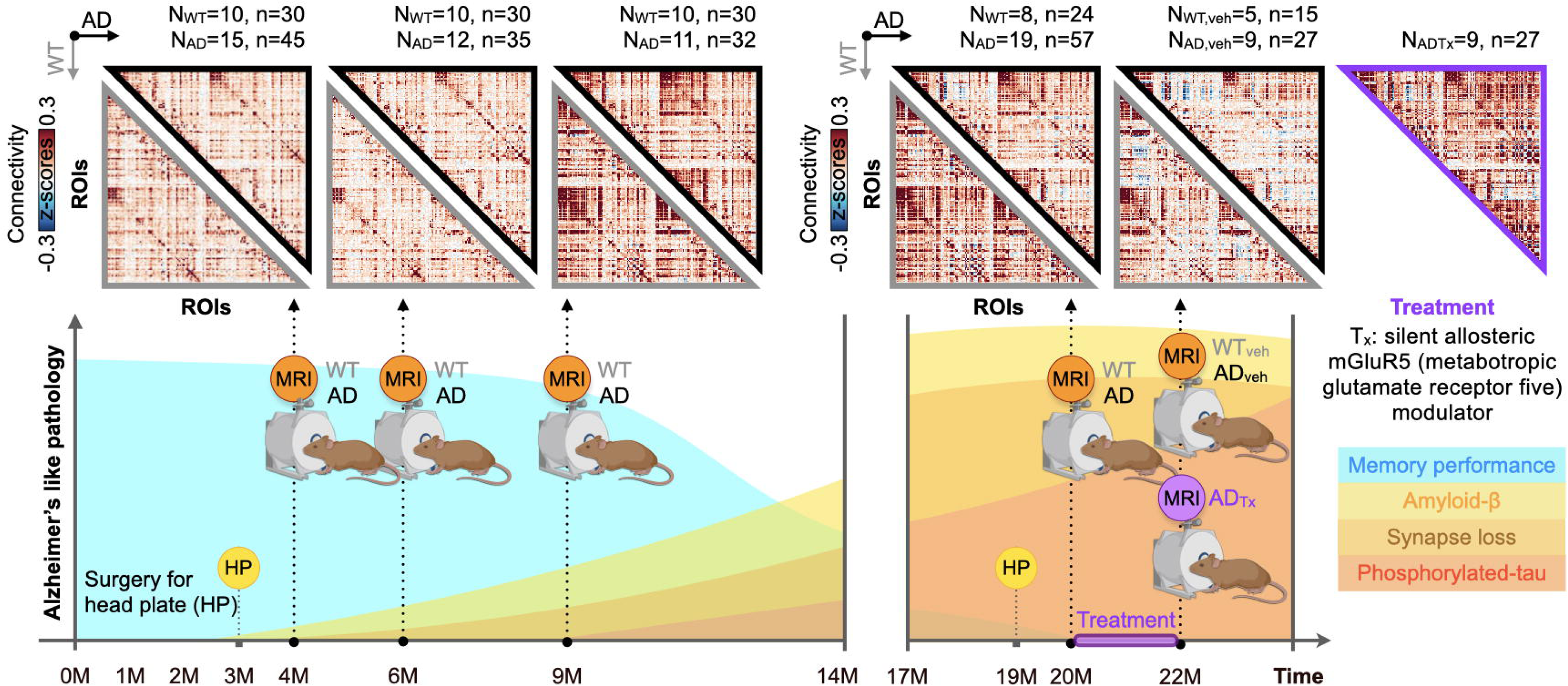
Schematic of AD-like pathology and study overview. Based on previous work in this AD-model,^26,27,29^ the approximate trajectory of AD-like pathology is illustrated on the timeline in warm colours: β-amyloid accumulation (yellow), synapse loss (orange) and phosphorylated-tau accumulation (red). Decreasing performance on memory tests (e.g., Y-maze test)^26^ is illustrated in blue. Imaging timepoints (MRI, orange) are indicated by black dotted arrows. A month prior to the first imaging timepoint (in each group, **Table 1**), all mice undergo a minimally invasive surgical procedure^34^ where a head-plate (HP) is affixed to the skull to reduce susceptibility artifacts and motion (HP, yellow). The treatment (Tx) period is indicated on the timeline (purple). Average Z-scored connectomes for each group at each timepoint are shown above the timeline. An atlas of 172 ROIs is used to generate the connectomes, 2 regions are removed for visualization purposes. Data from WT mice are outlined in grey (lower left). Data from AD (and AD_vehicle_) groups are outlined in black (upper right). Data from the AD_treated_ group (at 22M) are outlined in purple. The number of mice (‘N’) and runs (‘n’) are indicated above each connectome. ROIs: regions of interest. Animal breakdown: 4M: N_WT_=5/5 M/F; N_AD_=7/8 M/F; 6M: N_WT_=5/5 M/F; N_AD_=5/7 M/F; 9M: N_WT_=6/4 M/F; N_AD_=6/5 M/F; 20M: N_WT_=5/3 M/F; N_AD_=12/7 M/F; 22M: N_WT_=5/0 M/F; N_ADveh_=6/3 M/F; N_ADTx_=5/4 M/F.

### Age-related changes in functional brain organization diverge between WT and AD mice

AD progression occurs atop a background of age-related changes in behaviour and brain function.^44,45^ We examine age-related changes by quantifying differences between connectomes across timepoints: 4 vs. 6M (early), 4 vs. 9M (mid), 4 vs. 20M (late), and 4 vs. 22M (very late), within WT and AD groups (**Fig. 2**).

**Fig. 2.**
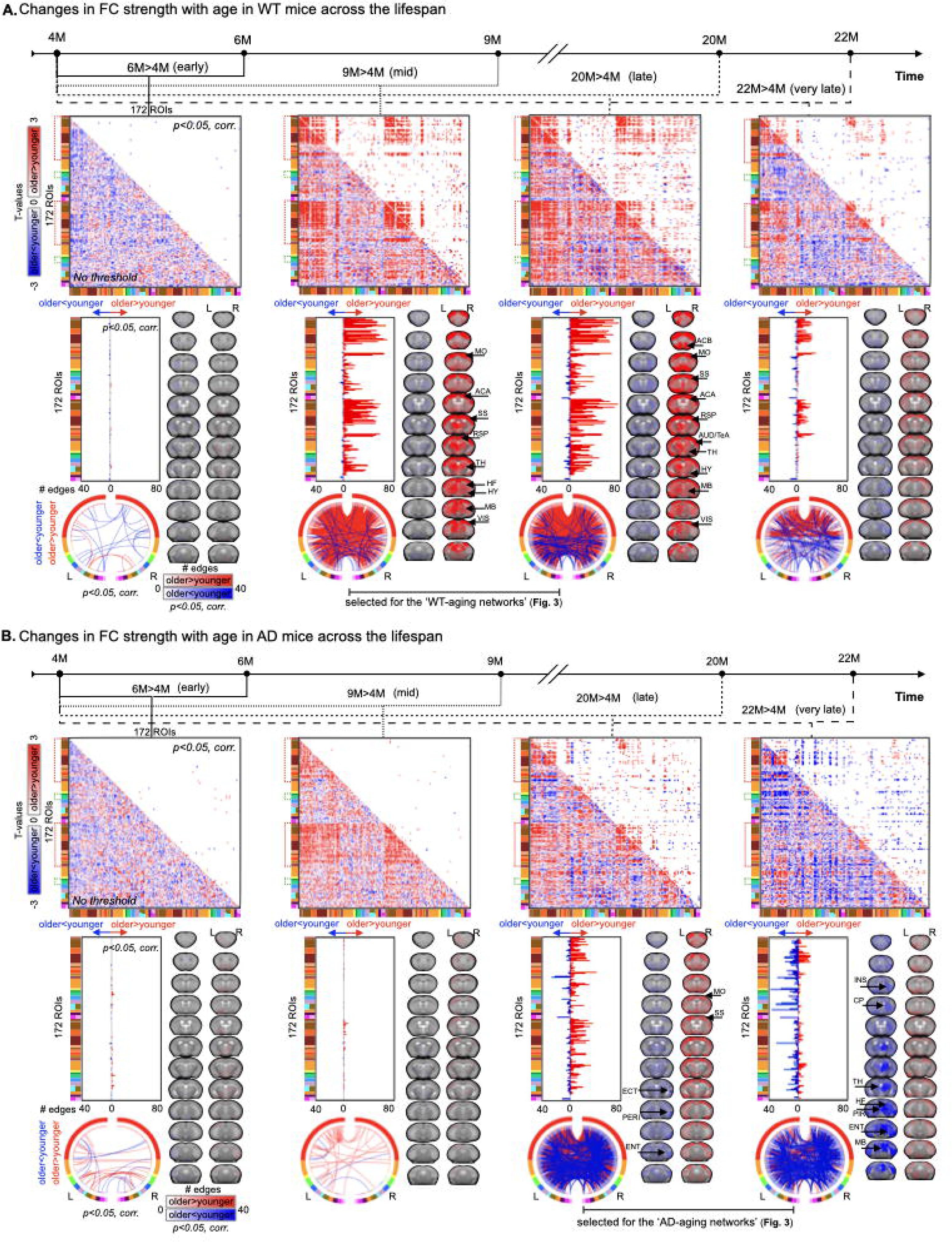
WT and AD follow distinct trajectories of age-related changes in functional brain organization. Two-sample t-tests are applied to uncover age-related changes in FC strength within WT and AD mice (**A.**, and **B.**, respectively). In both groups (WT and AD), the earliest timepoint (4M) is used as a baseline for the four subsequent imaging sessions. Thus, four stages of ‘aging’ are captured: early (6M vs. 4M), mid (9M vs. 4M), late (20M vs. 4M) and very late (22M vs. 4M). For connectomes, T-test results are displayed without (bottom half), and with (top half) a statistical threshold (corrected, *p<* 0.05). All other displays, vertical histograms of node degree, circle plots and node degree maps, results are shown after the statistical thresholds have been applied. For the brain region labels (rainbow) colour-scale, see **Fig. S1**. Positive t-values (red) indicate greater FC strength in older vs. younger mice. Negative values (blue) indicate the opposite. Node degree maps are displayed on anatomical reference images (greyscale). Clear and distinct aging effects are observed in both groups. The WT group shows an overall increase in FC strength in older vs. younger mice which emerges at 9M, persists at 20M, and shows some evidence of waning at 22M. Conversely, the AD group shows an overall decrease in FC strength in older vs. younger mice which emerges at 20M and persists until 22M. As a data reduction strategy, we summarize these data at the indicated timepoints for each group (Fig. 3).

Few early-life age-related changes in FC are identified in either group (**Fig. 2**, permutation test *p<*0.05). Beginning at mid-life, WT mice show a widespread increase in FC (**Fig. 2A**, permutation test *p<*0.001), that is not observed in AD mice (although, a trend in this direction is noticeable in unthresholded and uncorrected data, **Fig. 2B**, permutation test *p<*0.05). This increase, in the WT group, persists until the study endpoint. At 20 and 22M, some decreases in FC are also observed in ventral areas alongside more widespread increases (permutation test *p<*0.001). AD mice show a more modest FC increase with age, and this emerges later, in conjunction with a pronounced decrease in FC within subcortical regions (**Fig. 2B**). Overall, WT and AD groups follow distinct patterns of age-related changes in brain functional organization that unfold along different timescales. As a control, to account for any differences between WT and AD mice at 4M, we repeat our analyses using all mice (WT and AD) combined in a single group, at 4M, as a reference (**Fig. S2**). No differences in outcome patterns are observed.

Regions in WT mice that show pronounced age-related increases in FC beginning at 9M include somatosensory and motor regions (SS/MO), visual cortices (VIS), midbrain (MB), hypothalamus (HY) and the retrosplenial areas (RSP). AD mice show more modest increases in the same areas but starting at 20M. Moreover, AD mice show a more pronounced decrease in FC in the ventral parts of the brain encompassing the hippocampal formation (HF), entorhinal cortex (ENT), ectorhinal and perirhinal regions (ECT, PERI), ventral striatal areas including the caudoputamen (CP), and piriform and insular regions (PIR, INS).

### Aging trajectories in WT and AD mice effect overlapping and distinct networks

In addition to examining which regions are most affected by age, we can quantify over or under enrichment for edges within established or canonical networks. Here, and in following subsections, we use the triple-network model^33,46^: default mode, lateral cortical, and salience networks (DMN, LCN, and SN) as well as the edges that connect them (e.g., DMN to LCN) to characterize the data-driven brain circuits we identify. For a breakdown of the regions within these networks, see **Fig. S1B.** Above (**Fig. 2**), we establish that WT and AD mice show distinct patterns of age-related changes in FC. Here, we define a data-derived ‘WT-aging’ (**Fig. 3A**) and an ‘AD-aging’ (**Fig. 3B**) network. To do so, we select both the sum of *all* edges surviving correction (**Fig. 3A,B**, left), and the intersect of the *common* edges surviving correction (**Fig. 3A,B**, middle), at the two indicated timepoints. This will establish two sets of edges, a comprehensive (sum) and stringent (intersect) one, which will be used to define our ‘aging’ networks. Importantly, we collapse across the two timepoints that show the most pronounced effects: mid & late for WT and late & very late for AD (as indicated in **Fig. 2**). Note that the edges identified in both groups at the chosen stages are highly overlapping (**Fig. 3**, observed jaccard similarity significantly different from a null distribution, *p<*0.001, for both groups).

**Fig. 3.**
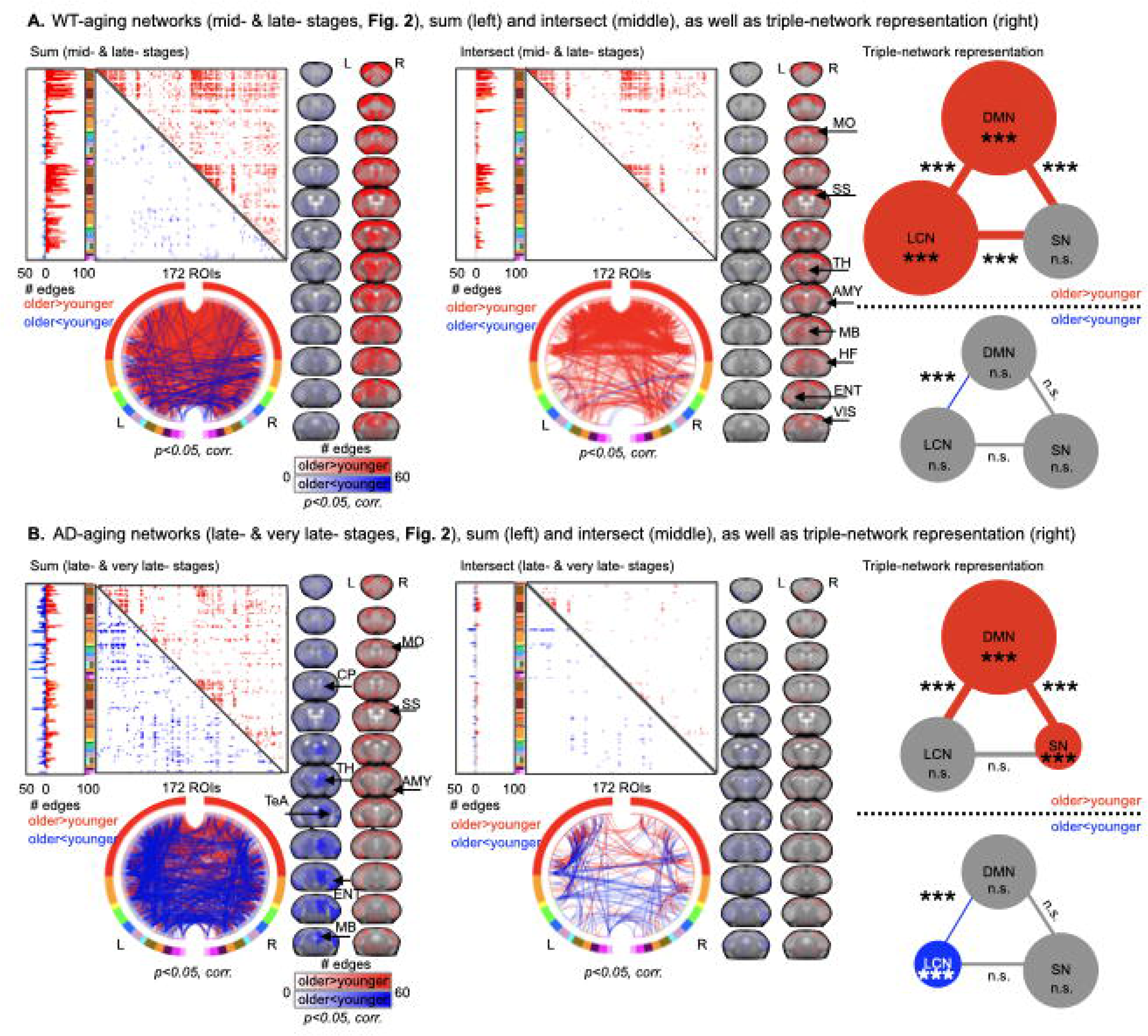
Summaries of data-derived WT- and AD-aging networks. We consider both the sum (all edges across two stages) and intersect (common edges across two stages) as a comprehensive (sum) and stringent (intersect) definition of our data-driven aging networks (left and middle panels respectively, **A.** and **B.**). Edges at chosen stages (Fig. 2) are highly overlapping (difference from null distribution, *p<*0.001) for both WT (**A.**) and AD (**B.**) groups. Aging networks are displayed as connectomes (172 x 172 ROIs, see **Fig. S1**). Greater FC in older relative to younger mice is shown in red (upper right in connectomes). Greater FC in younger relative to older mice is shown in blue (lower left in connectomes). Data are also shown as vertical histograms of node degree, circle plots, and node degree maps displayed on anatomical reference images (greyscale) in left and middle panels (**A.** and **B.**). All results have a statistical threshold (corrected, *p<*0.05) applied. Fisher’s exact test (MATLAB) is used to assess (under) enrichment for edges within data-derived networks and the triple-network model: DMN, LCN, and SN, as well as their inter-network pairs (corrected, permutation test x1,000 iterations). These results are depicted using ball-and-stick diagrams (right panel). An enlarged ball indicates enrichment (e.g., **A.** WT-aging DMN, older>younger), whereas a shrunken ball indicates under-enrichment (e.g., **A.** WT-aging SN, older>younger). Balls and sticks showing no (under) enrichment are drawn in grey (n.s.) whereas (under) enrichment is indicated by colour (red for older>younger, and blue for older<younger). Significance is displayed as * for *p<*0.05, ** for *p<*0.01 and *** for *p<*0.001. Equivalent findings are uncovered for triple network representation when we use the sum or intersect definition.

We compute enrichment (or depletion, under enrichment) for edges within the triple-network model and our aging networks (**Fig. 3A,B** right) using Fisher’s exact test as well as permutation testing (×1,000 iterations). Results are corrected for multiple comparisons. We find that the WT-aging positive (+ve) network (older>younger) is enriched for edges in the DMN and LCN as well as edges that connect all three networks. The WT-aging negative (−ve) network (older<younger) shows under-enrichment (depletion) for DMN to LCN. Further, the AD-aging (+ve) network (older>younger) shows enrichment in the DMN, DMN to LCN and DMN to SN, and under-enrichment in the SN. While the AD-aging (−ve) network (older<younger) shows under-enrichment in the LCN, DMN to LCN and a trend towards under-enrichment in the DMN. In sum, WT and AD mice show some common trends for which networks in the triple-network model are implicated in age-related changes in connectivity strength, but also some clear differences.

### Depressed connectivity strength in AD relative to WT mice emerges at 9M

After establishing aging trajectories for both genotypes (WT and AD), we compute differences between WT and AD connectomes at each timepoint (**Fig. 4**). Edges where FC in AD is greater than WT (AD>WT) are plotted as positive t-values (red). Edges where FC in AD<WT are plotted as negative t-values (blue). Permutation testing is applied (as above). Some very sparse but significant differences between WT and AD mice are observed at early timepoints (4 and 6M, *p<*0.05) with strong effects emerging by 9M (permutation test, *p<*0.001). Overwhelmingly, AD mice show lower FC. Notably, affected areas are bilaterally symmetric and appear throughout the brain but are concentrated in the cortex. Identified regions include the SS, MO, VIS, and RSP areas, as well as regions encompassing the HF, temporal association areas (TeA), nucleus accumbens (ACB), auditory (AUD) and MB areas.

**Fig. 4.**
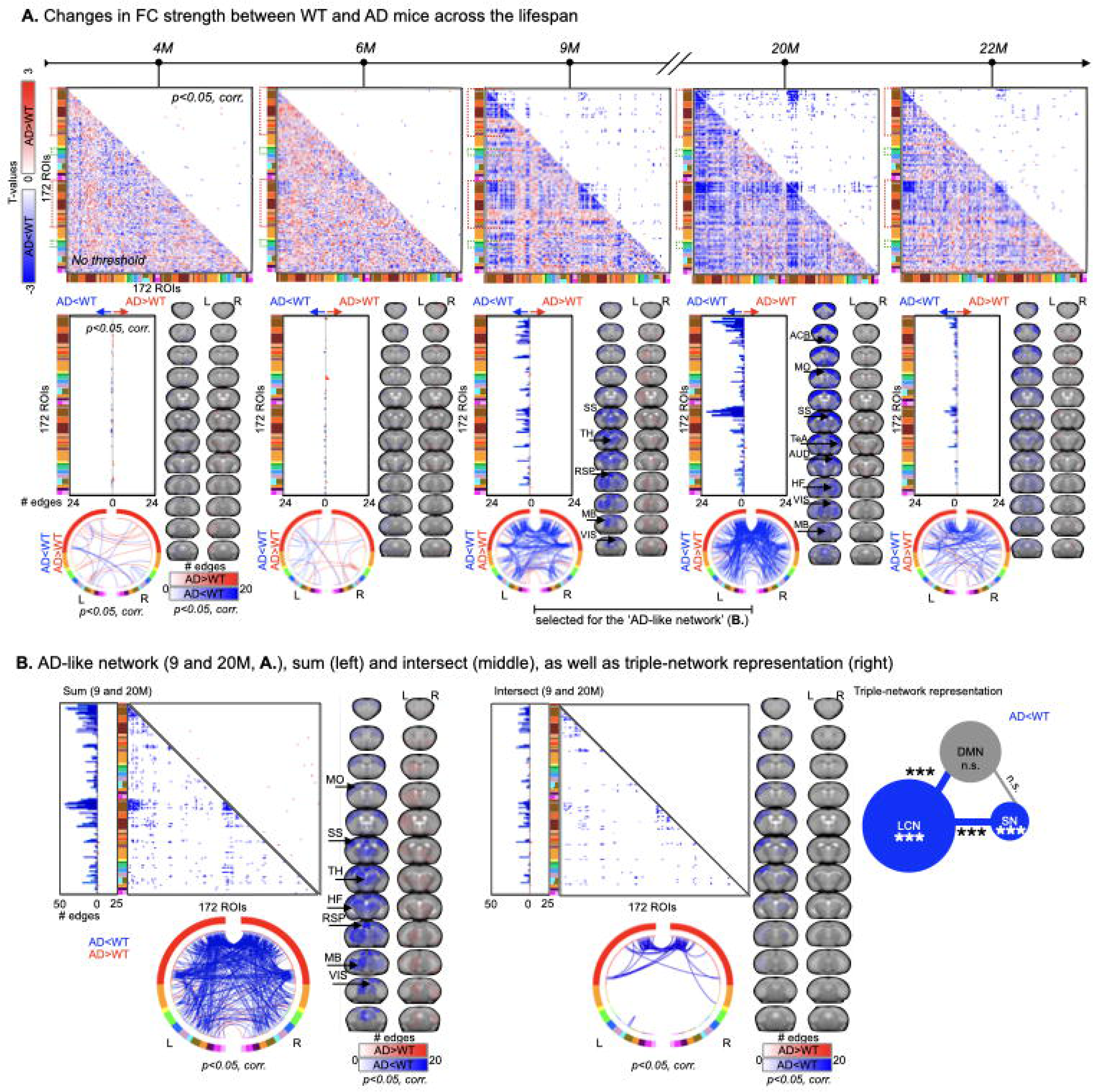
Emerging at 9M, AD mice show a pronounced decrease in FC strength relative to WT mice. (**A.**) As in Fig. 2 except here, we compare WT to AD groups at each timepoint. Positive t-values (red) indicate greater FC strength in AD vs. WT mice. Negative values (blue) indicate the opposite. (**B.**) As in Fig. 3 except here, we collapse across 9 and 20M timepoints (as indicated in **A.**). Significance is displayed as * for *p<*0.05, ** for *p<*0.01 and *** for *p<*0.001.

### The networks most strongly affected in AD relative to WT are the DMN and LCN

As above, we investigate the network composition^33,46^ of the brain circuits most affected in AD relative to WT mice. The most pronounced differences are observed at 9 and 20M (permutation test, *p<*0.001, **Fig. 4A**) and the edges identified at these timepoints are highly overlapping (observed jaccard similarity vs. null distribution, *p<*0.001). As above, we consider the sum (comprehensive) and intersect (stringent) in our definition of a data-derived ‘AD-like network’ (**Fig. 4B**, left/middle). We find that, at both thresholds, the AD-like (−ve) network (AD<WT) is enriched for edges in the DMN and LCN as well as edges that connect the DMN to the LCN, and LCN to SN (Fisher’s exact test; **Fig. 4B**, right). This network is also under-enriched for edges that are between the DMN and SN. A trend is noted for under enrichment for edges in the SN. There are no significant AD-like (+ve) network (AD>WT) findings.

### Comparison of mouse aging and AD networks with human literature meta-analysis (Neurosynth)

In an exploratory analysis, we use Neurosynth (http://neurosynth.org) to compare our mouse-model data-derived aging and AD-like networks to human neuroimaging studies using the queries ‘Age’ and ‘Alzheimer’ (**Fig. S3**). Neurosynth accepts one-word queries and provides ‘activation maps’ proportional to ‘hits’ in the literature without directionality (increases/decreases).^47^ As such, our aim is to visualize Age/Alzheimer maps across species for a purely qualitative comparison. **Fig. S3** shows the association test maps (z-scored, corrected for false discovery rate, FDR=0.01)^47^ from human fMRI data next to the node degree maps we obtain from mouse fMRI data. Encouragingly, in each comparison, homologous regions emerge. Specifically, somatomotor, SS, and dorsolateral-prefrontal areas are implicated in (healthy *and* WT) aging across species (**Fig. S3**, top), as are VIS, insula and RSP cortices (i.e., including RSP and precuneus in humans).^23^ In AD, subcortical areas, including the HF and TH, appear in both species (**Fig. S3**, bottom).

### Treatment with BMS-984923 rescues FC deficits in AD mice at 22M

After imaging at 20M, half of the mice in the AD group are randomly assigned to treatment with BMS-984923 (AD_treated_). The drug is incorporated into mouse chow to achieve sustained therapeutically effective levels, as documented in our previous study.^29^ We characterize treatment effects using three comparisons: (1) 22M WT vs. 22M AD_treated_ (**Fig. 5A**), (2) 22M AD_treated_ vs. 20M AD_vehicle_ (**Fig. 5B**, left), and (3) 22M AD_vehicle_ vs. 22M AD_treated_ (**Fig. 5B**, right). Across all comparisons we observe evidence of treatment effects. In comparison 1, there is a lack of FC differences between 22M WT and 22M AD_treated_ (**Fig. 5A** right, p>0.5 for the permutation test), even though there is a persistent difference between 22M WT and 22M AD_vehicle_ (**Fig. 5A**, left, reproduced from **Fig. 4**, *p<*0.001 for the permutation test). Comparisons 2 and 3 highlight edges whose strengths change with treatment. We select the brain circuits identified in comparisons 2 and 3 to generate, as before, a comprehensive and a stringent data-derived ‘treatment network’ (**Fig. 5C**, left/middle; summary of all data-derived networks in **Fig. S4**). Regions within this network include the VIS, AUD, MB, HY, HF, TH and CP areas. The (+ve) network (treated>vehicle) is under-enriched for edges in the LCN, and edges between the LCN and DMN (**Fig. 5C**, right). There are no significant (−ve) network (treated<vehicle) findings. Of note, the difference in FC at 22M between WT and AD_vehicle_ show a smaller effect size relative to the 20M timepoint. We find that this is likely due to a smaller sample size at the later timepoint by randomly selecting scans at 20M to match the number available at 22M and finding a similarly weakened effect size (data not shown).

**Fig. 5.**
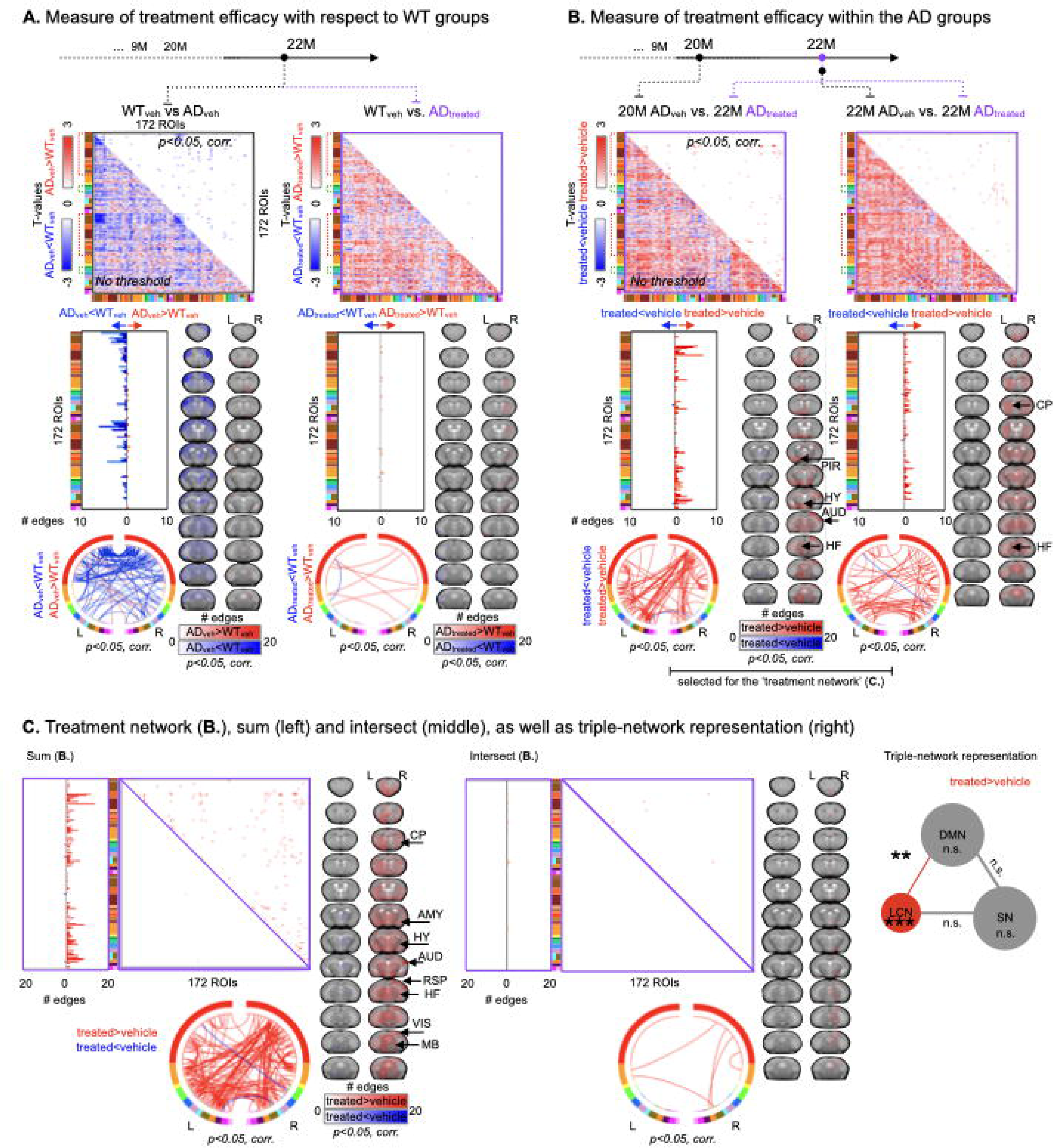
BMS-984923 treatment rescues FC deficits in AD_treated_ mice. (**A.**) Differences in FC strength between WT_vehicle_ and AD_vehicle_ (left, replicated from Fig. 4A) as well as WT_vehicle_ and AD_treated_ (right) at 22M. Positive t-values (red) indicate increased FC strength in AD_vehicle_ vs. WT mice (left) or increased FC strength in AD_treated_ vs. WT mice (right). Negative t-values (blue) indicated decreased FC strength in AD_vehicle_ vs. mice (left) or decreased FC strength in AD_treated_ vs. WT mice (right). BMS-984923 treatment appears to globally reduce FC strength differences between groups at 22M (two-sampled t-test, MATLAB, ttest2). (**B.**) Treatment effects in AD_treated_ (22M) relative to age-matched AD_vehicle_ mice (right) or AD_vehicle_ mice at 20M (left). Positive t-values (red) indicate increased FC in AD_treated_ vs. AD_vehicle_ mice. Negative t-values (blue) indicate decreased FC strength in AD_treated_ vs. AD_vehicle_ mice. (**C.**) As in Fig. 3 and Fig. 4B except here, we collapse across treatment effects (as indicated in **B.**). Significance is displayed as * for *p<*0.05, ** for *p<*0.01 and *** for *p<*0.001. VIS: visual cortex, AUD: auditory cortex, HF: hippocampal formation, HY: hypothalamus, AMY: amygdala, CP: caudoputamen.

Regions that show the highest degree of association with treatment include the VIS, MB, AUD, HY (**Fig. S5**). Notably, these regions also show high-degree representation in the AD-aging and AD-like networks (AD-aging and AD-like in **Fig. S5**) suggesting treatment specific targeting of regions involved in AD (as opposed to indirect or off-target effects of drug treatment).

### Mapping the shared functional landscape of data-derived networks

Seven data-derived networks are identified: AD-aging (−ve/+ve), WT-aging (−ve/+ve), AD-like (−ve/+ve) and treatment (+ve). We quantify the amount of cross-network (under) enrichment (as above), i.e., whether there is a significant number of shared edges, (or lack of shared edges), using Fisher’s exact test (and permutation testing, ×1,000 iteration), between all pairs of data-derived networks (**Fig. 6**). By definition, there is under-enrichment between AD-aging (+ve) vs. (−ve), and WT-aging (+ve) vs. (−ve) (**Fig. 6A,B**). We also find under-enrichment between AD-aging (+ve) vs. WT-aging (−ve), AD-aging (−ve) vs. WT-aging (+ve), and WT-aging (−ve) vs. AD-like (−ve). In addition, two patterns of enrichment emerge: the first between (+ve) aging (AD and WT) and AD-like (−ve), (**Fig. 6E**, left) and the second between treatment (+ve) and AD-aging (−ve), WT aging (+ve), and AD-like (−ve) (**Fig. 6E**, right). The second pattern is weaker, without as many edges, and shows no representation in the triple-network model (i.e., shared edges do not belong to DMN, LCN, SN or their inter-network pairings). Conversely, the first pattern shows a higher proportion of shared edges across network pairs (**Fig. 6A,E**, left). Edges shared between AD-aging (+ve) and AD-like (−ve) include 30% of the smaller, AD-like (−ve), network (proportion of shared edges, **Fig. 6A**, is computed as the fraction of the smaller network). Further, this 30% is fully contained within the other two inter-network pairings (**Fig. 6E**, left). The remaining pairings: WT-aging (+ve) vs. AD-aging (+ve) as well as WT-aging (+ve) vs. AD-like (−ve), show an even higher proportion of shared edges (67 and 85%, respectively, **Fig. 6A**), but a more modest inter-network pairing overlap (35%, **Fig. 6E**, left). Edges within the three pairings in the second pattern show a high representation of DMN and LCN, as well as edges that link DMN to LCN, and LCN to SN. Two of the three, AD-aging (+ve) vs. WT-aging (+ve) and AD-aging (+ve) vs. AD-like (−ve), show under-enrichment for the SN (**Fig. 6C**). Brain regions with high-degree that appear within these inter-network pairings (**Fig. 6D,F**) include primary SS, MO, VIS and RSP areas.

**Fig. 6.**
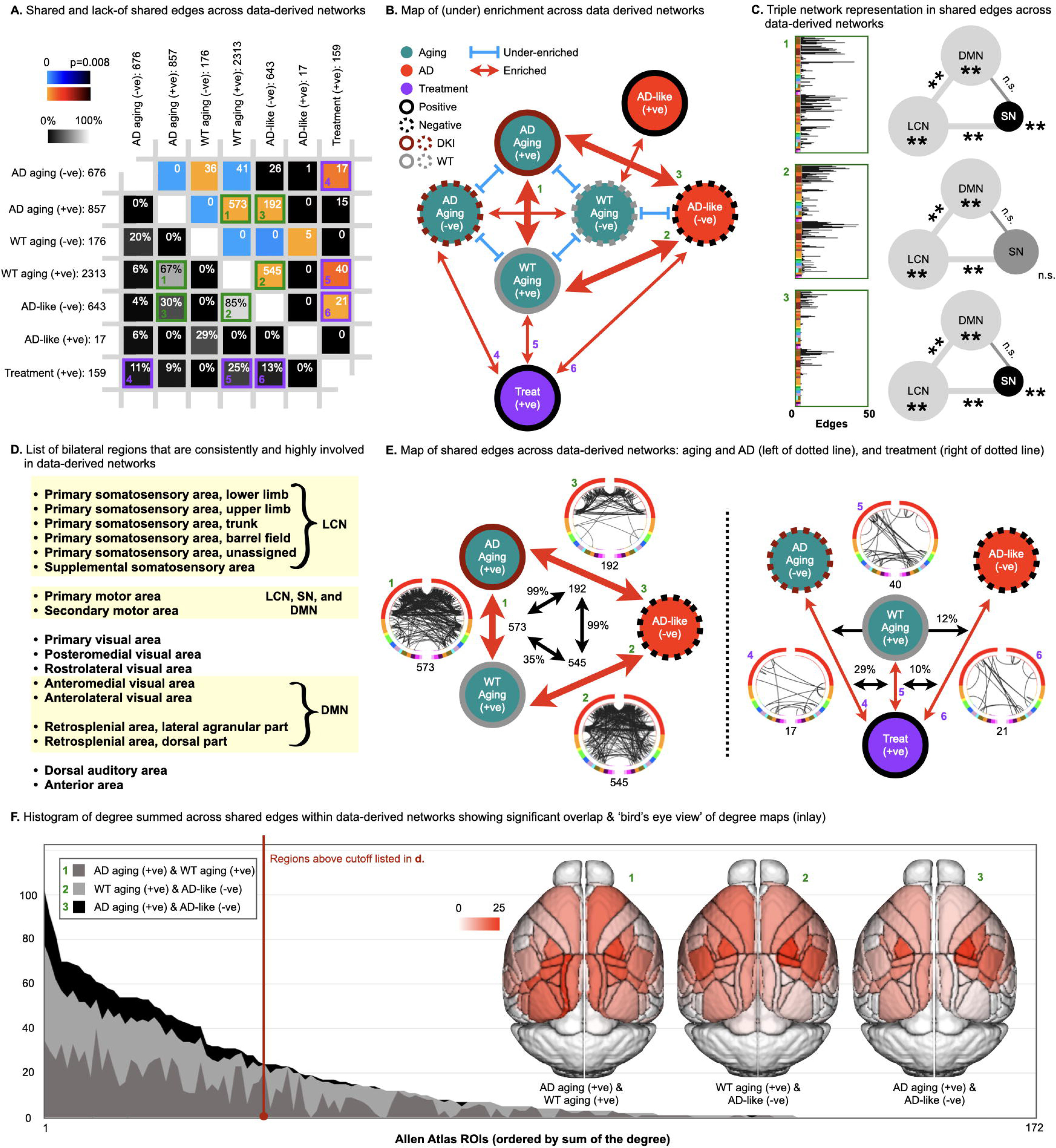
Shared functional anatomy of data-derived networks. (**A.**) Greyscale lower left: the fraction of shared edges between data-derived networks computed as the fraction of the smaller network contained within the larger network (e.g., 85% of the 643 edges within the AD-like (−ve) network are within the 2,313 edges within the WT-aging (+ve) network). Colour upper right: Using Fisher’s exact test (corrected, permutation test x1,000), we determine if a significant number of shared edges (hot colours), or a significant (*p<*0.008) lack of shared edges (cool colours), are found between data-derived network pairs. White numbers in each cell are the number of shared edges. (**B.**) Schematic of overlap (red arrows) and lack of overlap (blue lines with caps) between data-derived networks. Arrow size is proportional to significance. (**C.**) Edges shared between AD-aging (+ve), WT-aging (+ve), and AD-like (−ve) pairs – green: 1, 2, and 3 – are shown as vertical ROI plots and ball-and-stick diagrams. (**E.**) Schematic of overlap between two groupings of edges that are shared across data-derived networks: Age and AD (left of dotted line, green: 1, 2, and 3), and treatment (right of dotted line, purple: 4, 5, and 6). No significant representation of edges shared across data-derived networks including the treatment network – purple: 4, 5, and 6 – is found with the triple-network system (DMN, LCN, or SN). Bilateral regions that are identified consistently (in 1, 2, and 3) which show the highest degree are listed (**D.**). A layered histogram of degree for the 1, 2, and 3 grouping (**F.**) with a bird’s eye view of a degree colour-map (red), in-lay. Significance is displayed as * for *p<*0.05, ** for *p<*0.01 and *** for *p<*0.001.

### Mean FC within data-derived networks across the mouse lifespan

The product of each data-derived network and each mouse’s connectome is computed at each timepoint. To this end, each mouse’s connectome is masked using each of the data-derived networks (binary), so that edges not included in the networks are set to zero. The mean FC of nonzero elements are plotted for WT and AD mice (**Fig. 7**, **S6**). The key metric of disease is the AD-like (−ve) network (**Fig. 7A**). Mean FC in this network is decreased in AD relative to WT mice at 9M and 20M (*p<*0.01), but not at earlier timepoints. At 22 M, the AD_vehicle_ mice show decreased mean FC in the AD-like (−ve) network relative to WT and relative to AD_treated_ mice, but AD-treated are indistinguishable from WT. The treatment (+ve) network by definition shows increased mean FC in AD_treated_ mice relative to AD_vehicle_ at 22M (**Fig. 7B**). There is a significant treatment effect on the AD pathology at the *p<*0.001 level for the two conditions (pathology and treatment; ANOVA, [*F*(2,20)=10.48, *p*=0.0008]). Importantly, this drug-responsive network exhibits statistically significant decreases for AD mice relative to WT mice preceding treatment at 9M and 20M. The WT-aging (+ve) network associated with healthy brain maturation highlights an increase in mean FC in WT relative to AD mice at 9M and 20M (**Fig. S6**). Overall, connectivity in these data-driven networks across groups is well-aligned with the specific comparisons above.

**Fig. 7.**
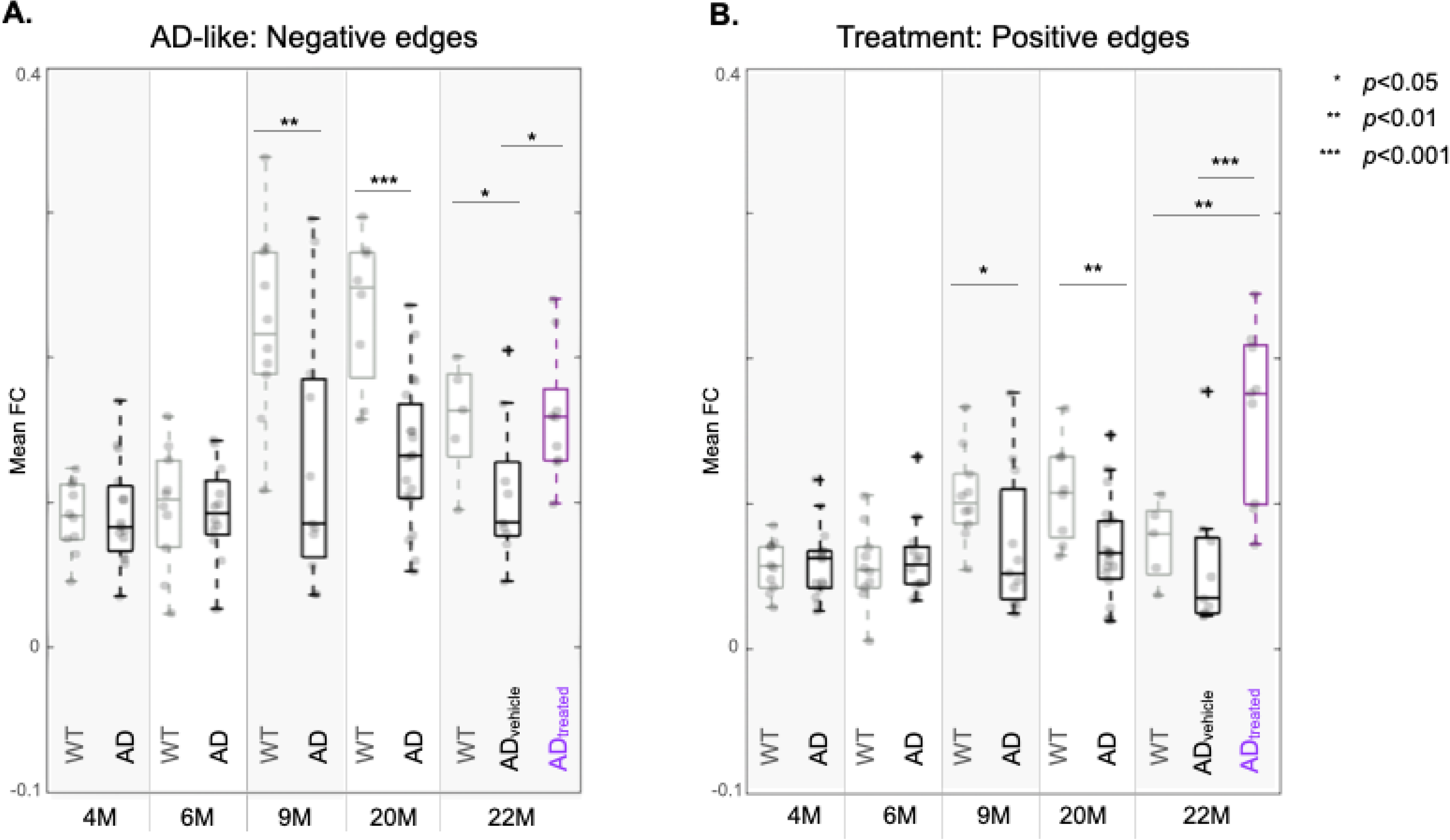
Mean FC within data-derived networks. Mean FC computed by multiplying data-derived networks by connectomes obtained from each mouse at each imaging timepoint. Boxplots are shown for WT (grey), and AD (vehicle/treated black/red) mice, for AD-like (−ve) (**A.**) and treatment (+ve) (**B.**) networks. The horizontal line indicates the median within each group. A t-test is used to compare mean FC strengths across groups, at each timepoint. For the 22M timepoint, an ANOVA is performed first, and t-tests are computed after to check the level of the significance found. Significance is displayed as * for *p<*0.05, ** for *p<*0.01, and *** for *p<*0.001.

## Discussion

Non-invasive imaging technologies, including fMRI, applied in animal models of human disease allow aspects of pathology to be characterized in a controlled environment, on a tractable timescale, and for the assessment of clinically accessible bioimaging markers of disease emergence, progression, and response to treatment. Here, we investigate the BOLD-fMRI correlates of AD, throughout the mouse lifespan, in an optimized model of disease.^29,48^ In addition, we investigate the neuroimaging correlates of a novel synapse rescuing treatment (BMS-984923) administered late in life, once AD-like pathology is well established. We uncover networks which characterize aging (with and without disease), disease-associated changes in FC strength, as well as treatment-response. Analyses are conducted using data-driven approaches (i.e., no *a priori* assumptions are made about the roles established brain circuits, e.g., DMN, or regions play in these processes). We quantify the presence (or absence) of classic large-scale networks following the discovery phase and examine correspondences across species.

We explore changes in FC strength with age in WT and AD mice at three life-stages: young-adulthood (6M or 20-30 years-of-age in humans), middle-age (9M or 38-47 years), and older-adulthood (20-22M or 56-69 years).^28^ We find FC strength in WT mice follows an inverted-U-shaped profile (**Fig. 2A, S6A**). This pattern is roughly in-line with a small pool of existing work using fMRI in mice^49,50^ (**Table 2**). Across studies, relative to baseline (2-4M), there is good agreement that circuit-specific increases in FC strength emerge and peak within 5-9M.^49,50^ Between 10-18M (which we do not examine), fMRI^49,50^ shows a return to baseline, and optical imaging methods^51^ show a dip below baseline (at 17-18M). At 20M, we observe a persistent increase in FC strength, followed by a reduction at 22M. Differences in these collective observations in later-life could be due to a mismatch between timepoints, modalities (optical vs. fMRI), anaesthesia (Egimendia et al.^50^ and Vasilkovska et al.^49^ use medetomidine and isoflurane whereas optical data in the Albertson et al.^51^ study is acquired from awake animals), or the specific brain circuits being examined. It is also possible that the inverted-U-shaped profile is incomplete, and the brain follows an even more complex double-humped shaped profile. Despite the small number of existing studies, increased FC strength in mid-life seems to be a relatively reproducible observation. Encouragingly, this inverted-U-shaped profile in fMRI FC strength is also reported in cognitively healthy human subjects. As in mice, middle-aged people show greater FC strength, relative to the young and elderly.^52,53^

**Table 2.** Change in FC strength with age in healthy mice.

| Age | This study | Egimendia | Vasilkovska | Albertson* |
| --- | --- | --- | --- | --- |
| 2-4M | Baseline | Baseline | Baseline | Baseline |
| 4-6M | • | • |  |  |
| 6-8M |  |  | ↑ |  |
| 8-10M | ↑ | ↑ |  |  |
| 10-12M |  |  | ↓ |  |
| 12-14M |  | ↓ |  |  |
| 14-16M |  |  |  |  |
| 16-18M |  |  |  | ↓ |
| 18-20M | ↑ |  |  |  |
| 20-22M | ↓ |  |  |  |

\* Optical imaging of Thy-1-GCaMP6f mice.

Further, in mice, we find good agreement across studies in terms of which brain regions show age-related changes in FC strength (**Table 3**). Universally, the DMN – which includes RSP and VIS areas among others – shows age-related changes.^52^ MO, ACA, as well as auditory and SS areas also show good-moderate cross-study agreement (given the methodological differences between studies). Notably, these implicated areas include regions that are not typically associated with cognition (e.g., SS and MO areas). We also note strong alignment across species. In humans, the DMN,^52,53^ as well as ‘cognitive’ and ‘motor’ circuits^54–56^ all show age-related changes in FC strength (that generally follow the inverted-U-shaped profile). In the present study, we also observe the LCN (includes SS, TeA, and MO areas), MB, HY, and TH to be involved in WT-aging. Given inter-study difference in how networks are defined, and analyses conducted (data-driven, vs. *a priori* network or ROI driven), it is unclear whether these findings are unique. Finally, our Neurosynth findings in the human brain associated with age (**Fig. S3**) show considerable overlap (including the TH and TeA) with the regions and networks identified in mice (**Table 3**).

**Table 3.** Brain areas showing changes in FC strength with age in WT-mice.

| Network/Region | This study | Egimendia | Vasilkovska | Albertson* |
| --- | --- | --- | --- | --- |
| DMN | • | • | • | • |
| RSP<sup>†</sup> | • | (part of DMN) | • | • |
| VIS<sup>†</sup> | • |  | • | • |
| MO<sup>†</sup> | • | • |  | • |
| AUD | (Fig. 6D) |  | • | • |
| ACA<sup>†</sup> | • |  | • | • |
| SS<sup>†</sup> | • | • |  |  |
| LCN | • |  |  |  |
| HY | • |  |  |  |
| TH<sup>†</sup> | • |  |  |  |
| ACB<sup>†</sup> | • |  |  |  |
| MB | • |  |  |  |
| PTL<sup>‡</sup> |  |  |  | • |

\* Optical imaging cannot access subsurface areas.
<sup>†</sup> Areas in our Neurosynth analysis (**Fig. S3**).
<sup>‡</sup> PTL: Posterior parietal association area.<sup>51</sup>
Refer to **Fig. S1** for abbreviations.

This emerging cross-study (and species) convergence is promising. Yet, there are several competing theories on the neurobiological drivers of healthy aging which have proven difficult to reconcile (see reviews)^57,58^ especially when using fMRI in isolation. Although it is the beyond the scope of the present work to interrogate the mechanisms which give rise to these fMRI-observable age-related changes, we are well positioned to interrogate their origins as part of our future work using multimodal approaches.^34,35,41^

Decreases in FC strength (AD vs. control) is an established hallmark of neuropathology in mouse models.^15,49,59,60^ Likewise, fMRI measures from humans show a tight association between cognitive decline and lower FC strength.^61,62^ Yet, these AD-related changes occur against a non-trivial background of aging that can be overlooked when a single timepoint is considered. Is lower FC strength (in AD vs. control) a result of a decrease in the AD group? Or (as our data suggest, **Fig. 2**, **6**, **7**, **S6**), a failure to increase with maturation? This is likely an important distinction that will require further investigation.

In addition to examining healthy aging, we examine age-related changes in FC strength in AD mice alongside differences between WT and AD groups at each timepoint. Changes in FC strength with age in AD mice (**Fig. 2**) emerge later in life (relative to WT aging) and show more widespread decreases (**Fig. 3**). Yet, there are also preserved common elements between WT and AD aging which are reflected by the high proportion of common circuitry (**Fig. 6**). From our inter-group (AD vs. WT) comparison (at each timepoint), we observe a clear decrease in FC strength by 9M in AD. The directionality of these FC changes (decrease in AD-like models compared to WT) is in-line with previous fMRI work in other mouse models of AD.^15–17,59^ Notably, the FC deficits reported here appear later in life (9M) compared to literature (as early as ~3M),^15^ which may be a result of the milder nature of the pathology in this mouse model. Critically, the DKI *App^NL-G-F^/hMapt* model^26,27^ investigated here has not previously been examined using fMRI. This is important, as this new model represents a step-forward in our ability to emulate AD-like pathology via its milder genetic profile.^63^

The circuits and regions we find to be involved in aging (**Fig. 2, 3**), WT vs. AD (**Fig. 4**), and their common elements (**Fig. 6**) are summarized in **Table 4**. Only two regions (ACB and HY) show age-related changes uniquely in WT mice, whereas more widespread age-related changes are unique to AD mice (INS, ECT, PERI, CP, and PIR). Regions that do not show age-related changes, but are identified when comparing WT vs. AD include TeA, PPA, ENT, and HF. Most circuits and regions, however, are shared across at least 2/3 comparisons (as expected based on our analyses, **Fig. 6**). Roughly, in order of frequency across comparisons (columns in **Table 4**), we observe involvement of the DMN-to-LCN and LCN (including SS and MO), the DMN (particularly VIS areas, as well as RSP, MO, and to a lesser extent ACA), TH, LCN-to-SN and MB. These findings, particularly the involvement of the DMN and LCN, are in-line with what has been observed in AD patients,^64,65^ individuals with mild cognitive impairment (MCI),^2,66^ the amyloid-positive but cognitively normal elderly,^7^ and healthy people of an advanced age.^67^

**Table 4.** Brain areas showing changes in FC strength across comparisons in this study.

| Network/Region |  | WT-Aging | AD-Aging | WT vs. AD | Shared (Fig. 6) |
| --- | --- | --- | --- | --- | --- |
| **DMN:** |  | ↑ (+ve); ↓ (-ve) | ↑ (+ve); ↓ (-ve) | • | ★ |
| VIS<sup>†</sup> |  | ↑ (+ve) | ↑ (-ve) | ↑ (-ve) | ★ |
| RSP<sup>†‡</sup> (PREC) |  | ↑ (+ve) | • | ↑ (-ve) | ★ |
| MO<sup>†</sup> |  | ↑ (+ve) | • | ↑ (-ve) | ★ |
| ACA<sup>†‡</sup> |  | ↑ (+ve) | • | ↑ (-ve) | • |
| **LCN:** |  | ↑ (+ve) | ↓ (-ve) | ↑ (-ve) | ★ |
| SS<sup>†</sup> |  | ↑ (+ve) | • | ↑ (-ve) | ★ |
| TeA |  | • | • | ↑ (-ve) | • |
| MO<sup>†</sup> |  | ↑ (+ve) | • | ↑ (-ve) | ★ |
| **SN:** |  | • | • | • | • |
| HY |  | ↑ (+ve) | • | • | • |
| TH<sup>†</sup> |  | ↑ (+ve) | ↑ (-ve) | ↑ (-ve) | • |
| INS<sup>†‡</sup> |  | • | ↑ (-ve) | • | • |
| MO<sup>†</sup> |  | ↑ (+ve) | • | ↑ (-ve) | ★ |
| ACA<sup>†‡</sup> |  | ↑ (+ve) | • | ↑ (-ve) | • |
| **DMN-LCN** |  | ↑ (+ve); ↓ (-ve) | ↑ (+ve); ↓ (-ve) | ↑ (-ve) | ★ |
| **DMN-SN** |  | ↑ (+ve) | ↑ (+ve) | • | • |
| **LCN-SN** |  | ↑ (+ve) | • | ↑ (-ve) | ★ |
| Other | MB | ↑ (+ve) | • | ↑ (-ve) | • |
|  | ECT | • | ↑ (-ve) | • | • |
|  | PERI | • | ↑ (-ve) | • | • |
|  | HF<sup>†</sup> | • | ↑ (-ve) | ↑ (-ve) | • |
|  | CP | • | ↑ (-ve) | • | • |
|  | PIR | • | ↑ (-ve) | • | • |
|  | PPA | • | • | ↑ (-ve) | • |
|  | ACB<sup>†</sup> | ↑ (+ve) | • | • | • |
|  | ENT | • | • | ↑ (-ve) | • |
|  | HF<sup>‡</sup> | • | • | ↑ (-ve) | • |
|  | AUD | • | • | • | ★ |
|  | AA | • | • | • | ★ |

ACB: Nucleus accumbens, ENT: entorhinal cortex.
<sup>†</sup>Neurosynth results for 'Age' (**Fig. S3**).
<sup>‡</sup>Neurosynth results for 'Alzheimer' (**Fig. S3**).
PREC: precuneus (human) to RSP (mouse).
Refer to **Fig. S1** for abbreviations.

Further, these patterns align (in time and space) with established neurobiological correlates of AD including Aβ-plaques^68^ and synapse losses^69^ across species. Specifically, the DMN shows *increased* FC in healthy adults where future preferential deposition of Aβ-plaques occur in (older) AD patients^70^ in parallel with *decreases* in FC.^64^ Further, regions in the DMN (including the RSP or PREC) are among the earliest to show both Aβ-pathology^65^ and change in FC in individuals with MCI.^66^ Mouse models show Aβ-pathology and synapse density changes in the DMN (RSP and VIS) and LCN (SS and MO), and HF.^26^ Mice also show alterations in FC that arise slightly before (at 6-9M) substantial increases in Aβ-pathology (which plateau – in the cortex – within the 8-10M window).^26^ Together, these findings hint at a relationship between changes in FC strength that *precede* Aβ-pathology and synapse loss, FC changes which position fMRI as a potentially useful tool for both AD identification and management.

In this regard, AD-associated cognitive deficits are attributable, at least in part, to synaptic failure triggered by Aβ and tau accumulation.^71,72^ Targeting a critical link in this cascade (see **Supplementary Discussion**), our group has shown that treatment with BMS-984923 elicits behavioural recovery, normalizes neuronal transcriptomics, prevents synaptic loss (if administered in early stages) and rescues synapse losses (when administered at late stages of the pathology, like in our study) in two mouse models of AD (inclusive of the DKI *App^NL-G-F^/hMapt*).^29,31^ Here, we conduct the first fMRI investigation of BMS-984923 treatment effects. To further characterize the therapeutic potential of this drug, we administer to older mice (19-9 vs. 20-22M), after AD-pathology is well established. In BMS-984923 AD_treated_ vs. WT mice (at 22M), we observe normalized FC (**Fig. 5A, S6A**). In AD_treated_ vs. AD_vehicle_ (at 20 or 22M) we observe significant increases in FC (**Fig. 5B**). Although a simple triple-network model is not emergent for these treatment-elicited changes in FC, regions highly involved in the salience network (TH, HY and CP) are detected. Additionally, there is significant overlap with areas implicated in our AD-aging (−ve), WT-aging (+ve), and AD-like (−ve) results (**Fig. 6E**, right).

Using FC in the identified AD-like (−ve) network as a disease measure (**Fig. 7**), the deficits observed in AD_vehicle_ relative to WT mice are fully corrected in AD_treated_ mice at 22 months after 2 months of intervention. Tracking connectivity in this network, or its human ortholog, provides a potential tool to monitor treatment benefit during translational development of this or other AD therapeutics with putative synapse rescuing benefit.

No study is without some limitations, and room for future work. Here, we use anaesthesia, the gold standard in rodent-fMRI, to minimize motion and stress.^43^ Yet, recent efforts to move the field towards performing experiments in awake mice are underway (see our own work in awake mice,^73^ and our recent systematic review).^74^ As discussed in detail within our review, there is still much we can learn from experiments in anesthetized animals, and from moving methodically towards performing experiments on awake animals. Further, this study does not include a group of WT_treated_ mice. This decision was made based on our previous work which showed no changes in synaptic density or single cell transcriptomics in a WT_treated_ cohort.^29^ Finally, although we include equal distributions of both biological sexes, we are not adequately powered for sex-based comparisons. This will be addressed as part of our future work.

In conclusion, we present a comprehensive investigation of the fMRI-FC correlates of AD in an optimized mouse model of the disease across the lifespan (4-22M). A detailed consideration of aging in both healthy-WT mice and their AD-counterparts is included alongside more classic groupwise comparisons. Overall, our findings align well with the existing literature from other mouse models and the human fMRI literature. We observe progressive AD-related dysconnectivity that emerges in known AD-vulnerable areas (sites of Aβ-accumulation, Tauopathy, and synaptic loss). Administration of a synapse-targeting treatment (BMS-984923) rescues the observed FC deficits in late-stage disease indicating the potential utility of this therapeutic as a synapse *rescuing* treatment and the role neuroimaging with fMRI could play in therapeutic development and patient management.

## Acknowledgments

Dr. Carolyn Fredericks for a close read of the manuscript and helpful feedback. Drs. Joel Greenwood, Omer Mano, and Paul Shamble from the Neurotechnology Core of the Yale Kavli Institute for Neuroscience for their technological expertise. Drs. Michael Crair and Ali Hamodi for help with optimizing and implementing transverse sinus injection experiments. Dr. Yonghyun Ha for his technical expertise and coil mending abilities. Dr. Corey Horien for critical discussion and coding advice. Kristin DeLuca for mouse husbandry.

## Author contributions

FM, SMS, and EMRL designed the study. FM carried out the surgical procedures, data acquisition, analyses and wrote the manuscript. GDG, DO, XS, and XP contributed code. AQ and JO implemented the skull stripping algorithm. BM and AO helped with animal and tissue handling. MC supervised GDG and edited the manuscript. SMS and EMRL supervised the study.

## Supplementary material

Supplementary material is available at *Brain* online

## References

1. Sheline YI, Morris JC, Snyder AZ, et al. APOE4 allele disrupts resting state fMRI connectivity in the absence of amyloid plaques or decreased CSF Abeta42. J Neurosci. Dec 15 2010;30(50):17035–40. doi:10.1523/JNEUROSCI.3987-10.2010

2. Wang J, Zuo X, Dai Z, et al. Disrupted functional brain connectome in individuals at risk for Alzheimer’s disease. Biol Psychiatry. Mar 1 2013;73(5):472–81. doi:10.1016/j.biopsych.2012.03.026

3. Kim J, Jeong M, Stiles WR, Choi HS. Neuroimaging Modalities in Alzheimer’s Disease: Diagnosis and Clinical Features. Int J Mol Sci. May 28 2022;23(11)doi:10.3390/ijms23116079

4. Marquez F, Yassa MA. Neuroimaging Biomarkers for Alzheimer’s Disease. Mol Neurodegener. Jun 7 2019;14(1):21. doi:10.1186/s13024-019-0325-5

5. Wang X, Huang W, Su L, et al. Neuroimaging advances regarding subjective cognitive decline in preclinical Alzheimer’s disease. Mol Neurodegener. Sep 22 2020;15(1):55. doi:10.1186/s13024-020-00395-3

6. Chetelat G. Multimodal Neuroimaging in Alzheimer’s Disease: Early Diagnosis, Physiopathological Mechanisms, and Impact of Lifestyle. J Alzheimers Dis. 2018;64(s1):S199–S211. doi:10.3233/JAD-179920

7. Sheline YI, Raichle ME, Snyder AZ, et al. Amyloid plaques disrupt resting state default mode network connectivity in cognitively normal elderly. Biol Psychiatry. Mar 15 2010;67(6):584–7. doi:10.1016/j.biopsych.2009.08.024

8. Morris JC, Storandt M, McKeel DW, Jr., et al. Cerebral amyloid deposition and diffuse plaques in “normal” aging: Evidence for presymptomatic and very mild Alzheimer’s disease. Neurology. Mar 1996;46(3):707–19. doi:10.1212/wnl.46.3.707

9. Buckner RL, Andrews-Hanna JR, Schacter DL. The brain’s default network: anatomy, function, and relevance to disease. Ann N Y Acad Sci. Mar 2008;1124:1–38. doi:10.1196/annals.1440.011

10. Scheff SW, Price DA, Schmitt FA, Mufson EJ. Hippocampal synaptic loss in early Alzheimer’s disease and mild cognitive impairment. Neurobiol Aging. Oct 2006;27(10):1372–84. doi:10.1016/j.neurobiolaging.2005.09.012

11. Terry RD, Masliah E, Salmon DP, et al. Physical basis of cognitive alterations in Alzheimer’s disease: synapse loss is the major correlate of cognitive impairment. Ann Neurol. Oct 1991;30(4):572–80. doi:10.1002/ana.410300410

12. DeKosky ST, Scheff SW. Synapse loss in frontal cortex biopsies in Alzheimer’s disease: correlation with cognitive severity. Ann Neurol. May 1990;27(5):457–64. doi:10.1002/ana.410270502

13. Sze CI, Troncoso JC, Kawas C, Mouton P, Price DL, Martin LJ. Loss of the presynaptic vesicle protein synaptophysin in hippocampus correlates with cognitive decline in Alzheimer disease. J Neuropathol Exp Neurol. Aug 1997;56(8):933–44. doi:10.1097/00005072-199708000-00011

14. Morrison JH, Baxter MG. The ageing cortical synapse: hallmarks and implications for cognitive decline. Nat Rev Neurosci. Mar 7 2012;13(4):240–50. doi:10.1038/nrn3200

15. Mandino F, Yeow LY, Bi RZ, et al. The lateral entorhinal cortex is a hub for local and global dysfunction in early Alzheimer’s disease states. J Cerebr Blood F Met. Apr 25 2022;doi:Artn 0271678x221082016 10.1177/0271678x221082016

16. Grandjean J, Derungs R, Kulic L, et al. Complex interplay between brain function and structure during cerebral amyloidosis in APP transgenic mouse strains revealed by multi-parametric MRI comparison. Neuroimage. Jul 1 2016;134:1–11. doi:10.1016/j.neuroimage.2016.03.042

17. Shah D, Jonckers E, Praet J, et al. Resting state FMRI reveals diminished functional connectivity in a mouse model of amyloidosis. PLoS One. 2013;8(12):e84241. doi:10.1371/journal.pone.0084241

18. Mandino F, Cerri DH, Garin CM, et al. Animal Functional Magnetic Resonance Imaging: Trends and Path Toward Standardization. Front Neuroinform. Jan 22 2020;13 doi:ARTN 78 10.3389/fninf.2019.00078

19. Markicevic M, Savvateev I, Grimm C, Zerbi V. Emerging imaging methods to study whole-brain function in rodent models. Transl Psychiatry. Sep 4 2021;11(1):457. doi:10.1038/s41398-021-01575-5

20. Gorges M, Roselli F, Muller HP, Ludolph AC, Rasche V, Kassubek J. Functional Connectivity Mapping in the Animal Model: Principles and Applications of Resting-State fMRI. Front Neurol. 2017;8:200. doi:10.3389/fneur.2017.00200

21. Zerbi V, Grandjean J, Rudin M, Wenderoth N. Mapping the mouse brain with rs-fMRI: An optimized pipeline for functional network identification. Neuroimage. Dec 2015;123:11–21. doi:10.1016/j.neuroimage.2015.07.090

22. Grandjean J, Schroeter A, Batata I, Rudin M. Optimization of anesthesia protocol for resting-state fMRI in mice based on differential effects of anesthetics on functional connectivity patterns. Neuroimage. Nov 15 2014;102:838–847. doi:10.1016/j.neuroimage.2014.08.043

23. Xu N, LaGrow TJ, Anumba N, et al. Functional Connectivity of the Brain Across Rodents and Humans. Front Neurosci. 2022;16:816331. doi:10.3389/fnins.2022.816331

24. Gozzi A, Zerbi V. Modeling Brain Dysconnectivity in Rodents. Biol Psychiatry. Mar 1 2023;93(5):419–429. doi:10.1016/j.biopsych.2022.09.008

25. Sharma H, Chang KA, Hulme J, An SSA. Mammalian Models in Alzheimer’s Research: An Update. Cells. Oct 16 2023;12(20) doi:10.3390/cells12202459

26. Saito T, Matsuba Y, Mihira N, et al. Single App knock-in mouse models of Alzheimer’s disease. Nat Neurosci. May 2014;17(5):661–3. doi:10.1038/nn.3697

27. Saito T, Mihira N, Matsuba Y, et al. Humanization of the entire murine Mapt gene provides a murine model of pathological human tau propagation. J Biol Chem. Aug 23 2019;294(34):12754–12765. doi:10.1074/jbc.RA119.009487

28. Dutta S, Sengupta P. Men and mice: Relating their ages. Life Sci. May 1 2016;152:244–8. doi:10.1016/j.lfs.2015.10.025

29. Spurrier J, Nicholson L, Fang XT, et al. Reversal of synapse loss in Alzheimer mouse models by targeting mGluR5 to prevent synaptic tagging by C1Q. Sci Transl Med. Jun 2022;14(647):eabi8593. doi:10.1126/scitranslmed.abi8593

30. Huang LK, Kuan YC, Lin HW, Hu CJ. Clinical trials of new drugs for Alzheimer disease: a 2020-2023 update. J Biomed Sci. Oct 2 2023;30(1):83. doi:10.1186/s12929-023-00976-6

31. Haas LT, Salazar SV, Smith LM, et al. Silent Allosteric Modulation of mGluR5 Maintains Glutamate Signaling while Rescuing Alzheimer’s Mouse Phenotypes. Cell Rep. Jul 5 2017;20(1):76–88. doi:10.1016/j.celrep.2017.06.023

32. Menon V. Large-scale brain networks and psychopathology: a unifying triple network model. Trends Cogn Sci. Oct 2011;15(10):483–506. doi:10.1016/j.tics.2011.08.003

33. Mandino F, Vrooman RM, Foo HE, et al. A triple-network organization for the mouse brain. Mol Psychiatr. Feb 2022;27(2):865–872. doi:10.1038/s41380-021-01298-5

34. Lake EMR, Ge X, Shen X, et al. Simultaneous cortex-wide fluorescence Ca(2+) imaging and whole-brain fMRI. Nat Methods. Jul 2020;18(7):835. doi:10.1038/s41592-021-01217-0

35. Vafaii H, Mandino F, Desrosiers-Gregoire G, et al. Multimodal measures of spontaneous brain activity reveal both common and divergent patterns of cortical functional organization. Nat Commun. Jan 3 2024;15(1):229. doi:10.1038/s41467-023-44363-z

36. Desrosiers-Gregoire G, Devenyi GA, Grandjean J, Chakravarty MM. Rodent Automated Bold Improvement of EPI Sequences (RABIES): A standardized image processing and data quality platform for rodent fMRI. bioRxiv. 2022:2022.08.20.504597.

37. Manjon JV, Coupe P, Marti-Bonmati L, Collins DL, Robles M. Adaptive Non-Local Means Denoising of MR Images With Spatially Varying Noise Levels. J Magn Reson Imaging. Jan 2010;31(1):192–203. doi:10.1002/jmri.22003

38. Sled JG, Zijdenbos AP, Evans AC. A nonparametric method for automatic correction of intensity nonuniformity in MRI data. IEEE Trans Med Imaging. Feb 1998;17(1):87–97. doi:10.1109/42.668698

39. Atlas AMB. Allen institute for brain science. Available online at: mouse brain-map org. 2004;

40. Wang Q, Ding SL, Li Y, et al. The Allen Mouse Brain Common Coordinate Framework: A 3D Reference Atlas. Cell. May 14 2020;181(4):936–953 e20. doi:10.1016/j.cell.2020.04.007

41. O’Connor D, Mandino F, Shen X, et al. Functional network properties derived from wide-field calcium imaging differ with wakefulness and across cell type. NeuroImage. 2022;264:119735.

42. Grandjean J, Desrosiers-Gregoire G, Anckaerts C, et al. A consensus protocol for functional connectivity analysis in the rat brain. Nature neuroscience. 2023;26(4):673–681.

43. Grandjean J, Canella C, Anckaerts C, et al. Common functional networks in the mouse brain revealed by multi-centre resting-state fMRI analysis. Neuroimage. 2020;205:116278.

44. Bondi MW, Houston WS, Salmon DP, et al. Neuropsychological deficits associated with Alzheimer’s disease in the very-old: discrepancies in raw vs. standardized scores. J Int Neuropsychol Soc. Jul 2003;9(5):783–95. doi:10.1017/S1355617703950119

45. Betzel RF, Byrge L, He Y, Goni J, Zuo XN, Sporns O. Changes in structural and functional connectivity among resting-state networks across the human lifespan. Neuroimage. Nov 15 2014;102 Pt 2:345–57. doi:10.1016/j.neuroimage.2014.07.067

46. Whitesell JD, Liska A, Coletta L, et al. Regional, Layer, and Cell-Type-Specific Connectivity of the Mouse Default Mode Network. Neuron. Feb 3 2021;109(3)doi:10.1016/j.neuron.2020.11.011

47. Yarkoni T, Poldrack RA, Nichols TE, Van Essen DC, Wager TD. Large-scale automated synthesis of human functional neuroimaging data. Nat Methods. Jun 26 2011;8(8):665–70. doi:10.1038/nmeth.1635

48. Zhang H, Wu L, Pchitskaya E, et al. Neuronal Store-Operated Calcium Entry and Mushroom Spine Loss in Amyloid Precursor Protein Knock-In Mouse Model of Alzheimer’s Disease. J Neurosci. Sep 30 2015;35(39):13275–86. doi:10.1523/JNEUROSCI.1034-15.2015

49. Vasilkovska T, Adhikari MH, Van Audekerke J, et al. Resting-state fMRI reveals longitudinal alterations in brain network connectivity in the zQ175DN mouse model of Huntington’s disease. Neurobiol Dis. Jun 1 2023;181:106095. doi:10.1016/j.nbd.2023.106095

50. Egimendia A, Minassian A, Diedenhofen M, Wiedermann D, Ramos-Cabrer P, Hoehn M. Aging Reduces the Functional Brain Networks Strength-a Resting State fMRI Study of Healthy Mouse Brain. Front Aging Neurosci. 2019;11:277. doi:10.3389/fnagi.2019.00277

51. Albertson AJ, Landsness EC, Tang MJ, et al. Normal aging in mice is associated with a global reduction in cortical spectral power and network-specific declines in functional connectivity. Neuroimage. Aug 15 2022;257:119287. doi:10.1016/j.neuroimage.2022.119287

52. Mak LE, Minuzzi L, MacQueen G, Hall G, Kennedy SH, Milev R. The Default Mode Network in Healthy Individuals: A Systematic Review and Meta-Analysis. Brain Connect. Feb 2017;7(1):25–33. doi:10.1089/brain.2016.0438

53. Staffaroni AM, Brown JA, Casaletto KB, et al. The Longitudinal Trajectory of Default Mode Network Connectivity in Healthy Older Adults Varies As a Function of Age and Is Associated with Changes in Episodic Memory and Processing Speed. J Neurosci. Mar 14 2018;38(11):2809–2817. doi:10.1523/JNEUROSCI.3067-17.2018

54. Bo J, Lee CM, Kwak Y, et al. Lifespan differences in cortico-striatal resting state connectivity. Brain Connect. Apr 2014;4(3):166–80. doi:10.1089/brain.2013.0155

55. Jolles DD, van Buchem MA, Crone EA, Rombouts SA. A comprehensive study of whole-brain functional connectivity in children and young adults. Cereb Cortex. Feb 2011;21(2):385–91. doi:10.1093/cercor/bhq104

56. Li Z, Petersen IT, Wang L, Radua J, Yang G, Liu X. Inverted U-shaped brain development of cognitive control across the human lifespan. bioRxiv. Aug 21 2023;doi:10.1101/2023.08.20.554018

57. McDonough IM, Nolin SA, Visscher KM. 25 years of neurocognitive aging theories: What have we learned? Front Aging Neurosci. 2022;14:1002096. doi:10.3389/fnagi.2022.1002096

58. Sala-Llonch R, Bartres-Faz D, Junque C. Reorganization of brain networks in aging: a review of functional connectivity studies. Front Psychol. 2015;6:663. doi:10.3389/fpsyg.2015.00663

59. Shah D, Praet J, Latif Hernandez A, et al. Early pathologic amyloid induces hypersynchrony of BOLD resting-state networks in transgenic mice and provides an early therapeutic window before amyloid plaque deposition. Alzheimers Dement. Sep 2016;12(9):964–976. doi:10.1016/j.jalz.2016.03.010

60. Zerbi V, Wiesmann M, Emmerzaal TL, et al. Resting-state functional connectivity changes in aging apoE4 and apoE-KO mice. J Neurosci. Oct 15 2014;34(42):13963–75. doi:10.1523/JNEUROSCI.0684-14.2014

61. Damoiseaux JS, Beckmann CF, Arigita EJ, et al. Reduced resting-state brain activity in the “default network” in normal aging. Cereb Cortex. Aug 2008;18(8):1856–64. doi:10.1093/cercor/bhm207

62. Onoda K, Ishihara M, Yamaguchi S. Decreased functional connectivity by aging is associated with cognitive decline. J Cogn Neurosci. Nov 2012;24(11):2186–98. doi:10.1162/jocn_a_00269

63. Sasaguri H, Hashimoto S, Watamura N, et al. Recent Advances in the Modeling of Alzheimer’s Disease. Front Neurosci. 2022;16:807473. doi:10.3389/fnins.2022.807473

64. Greicius MD, Srivastava G, Reiss AL, Menon V. Default-mode network activity distinguishes Alzheimer’s disease from healthy aging: evidence from functional MRI. Proc Natl Acad Sci U S A. Mar 30 2004;101(13):4637–42. doi:10.1073/pnas.0308627101

65. Sheline YI, Raichle ME. Resting state functional connectivity in preclinical Alzheimer’s disease. Biol Psychiatry. Sep 1 2013;74(5):340–7. doi:10.1016/j.biopsych.2012.11.028

66. Vann SD, Aggleton JP, Maguire EA. What does the retrosplenial cortex do? Nature Reviews Neuroscience. 2009/11/01 2009;10(11):792–802. doi:10.1038/nrn2733

67. Andrews-Hanna JR, Snyder AZ, Vincent JL, et al. Disruption of large-scale brain systems in advanced aging. Neuron. Dec 6 2007;56(5):924–35. doi:10.1016/j.neuron.2007.10.038

68. Hampel H, Hu Y, Hardy J, et al. The amyloid-beta pathway in Alzheimer’s disease: a plain language summary. Neurodegener Dis Manag. Jun 2023;13(3):141–149. doi:10.2217/nmt-2022-0037

69. Zhang J, Wang J, Xu X, et al. In vivo synaptic density loss correlates with impaired functional and related structural connectivity in Alzheimer’s disease. J Cereb Blood Flow Metab. Jun 2023;43(6):977–988. doi:10.1177/0271678X231153730

70. Buckner RL, Snyder AZ, Shannon BJ, et al. Molecular, structural, and functional characterization of Alzheimer’s disease: evidence for a relationship between default activity, amyloid, and memory. J Neurosci. Aug 24 2005;25(34):7709–17. doi:10.1523/JNEUROSCI.2177-05.2005

71. Selkoe DJ. Alzheimer’s disease is a synaptic failure. Science. Oct 25 2002;298(5594):789–91. doi:10.1126/science.1074069

72. Brier MR, Thomas JB, Ances BM. Network dysfunction in Alzheimer’s disease: refining the disconnection hypothesis. Brain Connect. Jun 2014;4(5):299–311. doi:10.1089/brain.2014.0236

73. Mandino F, Shen X, O’Connor D, et al. Longitudinal simultaneous cortex-wide Ca2+ imaging and whole-brain functional magnetic resonance imaging in awake mice. SAGE PUBLICATIONS INC 2455 TELLER RD, THOUSAND OAKS, CA 91320 USA; 2022:75–75.

74. Mandino F, Vujic S, Grandjean J, Lake EMR. Where do we stand on fMRI in awake mice? Cereb Cortex. Dec 13 2023;doi:10.1093/cercor/bhad478

75. Wang SJ, Peterson DJ, Gatenby JC, Li WB, Grabowski TJ, Madhyastha TM. Evaluation of Field Map and Nonlinear Registration Methods for Correction of Susceptibility Artifacts in Diffusion MRI. Front Neuroinform. Feb 21 2017;11 doi:ARTN 17 10.3389/fninf.2017.00017

76. Huang H, Degnan AP, Balakrishnan A, et al. Oxazolidinone-based allosteric modulators of mGluR5: Defining molecular switches to create a pharmacological tool box. Bioorg Med Chem Lett. Sep 1 2016;26(17):4165–9. doi:10.1016/j.bmcl.2016.07.065

77. Bachmanov AA, Reed DR, Beauchamp GK, Tordoff MG. Food intake, water intake, and drinking spout side preference of 28 mouse strains. Behav Genet. Nov 2002;32(6):435–43. doi:10.1023/a:1020884312053

78. Menon V. The Triple Network Model, Insight, and Large-Scale Brain Organization in Autism. Biol Psychiatry. Aug 15 2018;84(4):236–238. doi:10.1016/j.biopsych.2018.06.012

79. Lauren J, Gimbel DA, Nygaard HB, Gilbert JW, Strittmatter SM. Cellular prion protein mediates impairment of synaptic plasticity by amyloid-beta oligomers. Nature. Feb 26 2009;457(7233):1128–32. doi:10.1038/nature07761

80. Kostylev MA, Kaufman AC, Nygaard HB, et al. Prion-Protein-interacting Amyloid-beta Oligomers of High Molecular Weight Are Tightly Correlated with Memory Impairment in Multiple Alzheimer Mouse Models. J Biol Chem. Jul 10 2015;290(28):17415–38. doi:10.1074/jbc.M115.643577

81. Kostylev MA, Tuttle MD, Lee S, et al. Liquid and Hydrogel Phases of PrP(C) Linked to Conformation Shifts and Triggered by Alzheimer’s Amyloid-beta Oligomers. Mol Cell. Nov 1 2018;72(3):426–443 e12. doi:10.1016/j.molcel.2018.10.009

82. Um JW, Nygaard HB, Heiss JK, et al. Alzheimer amyloid-beta oligomer bound to postsynaptic prion protein activates Fyn to impair neurons. Nat Neurosci. Sep 2012;15(9):1227–35. doi:10.1038/nn.3178

83. Smith LM, Strittmatter SM. Binding Sites for Amyloid-beta Oligomers and Synaptic Toxicity. Cold Spring Harb Perspect Med. May 1 2017;7(5) doi:10.1101/cshperspect.a024075

84. Chen S, Yadav SP, Surewicz WK. Interaction between human prion protein and amyloid-beta (Abeta) oligomers: role OF N-terminal residues. J Biol Chem. Aug 20 2010;285(34):26377–83. doi:10.1074/jbc.M110.145516

85. Haas LT, Kostylev MA, Strittmatter SM. Therapeutic molecules and endogenous ligands regulate the interaction between brain cellular prion protein (PrPC) and metabotropic glutamate receptor 5 (mGluR5). J Biol Chem. Oct 10 2014;289(41):28460–77. doi:10.1074/jbc.M114.584342

86. Fluharty BR, Biasini E, Stravalaci M, et al. An N-terminal fragment of the prion protein binds to amyloid-beta oligomers and inhibits their neurotoxicity in vivo. J Biol Chem. Mar 15 2013;288(11):7857–7866. doi:10.1074/jbc.M112.423954

87. Haas LT, Strittmatter SM. Oligomers of Amyloid beta Prevent Physiological Activation of the Cellular Prion Protein-Metabotropic Glutamate Receptor 5 Complex by Glutamate in Alzheimer Disease. J Biol Chem. Aug 12 2016;291(33):17112–21. doi:10.1074/jbc.M116.720664

88. Salazar SV, Gallardo C, Kaufman AC, et al. Conditional Deletion of Prnp Rescues Behavioral and Synaptic Deficits after Disease Onset in Transgenic Alzheimer’s Disease. J Neurosci. Sep 20 2017;37(38):9207–9221. doi:10.1523/JNEUROSCI.0722-17.2017

89. Um JW, Kaufman AC, Kostylev M, et al. Metabotropic glutamate receptor 5 is a coreceptor for Alzheimer abeta oligomer bound to cellular prion protein. Neuron. Sep 4 2013;79(5):887–902. doi:10.1016/j.neuron.2013.06.036

90. Beraldo FH, Ostapchenko VG, Caetano FA, et al. Regulation of Amyloid beta Oligomer Binding to Neurons and Neurotoxicity by the Prion Protein-mGluR5 Complex. J Biol Chem. Oct 14 2016;291(42):21945–21955. doi:10.1074/jbc.M116.738286

91. Hamilton A, Esseltine JL, DeVries RA, Cregan SP, Ferguson SS. Metabotropic glutamate receptor 5 knockout reduces cognitive impairment and pathogenesis in a mouse model of Alzheimer’s disease. Mol Brain. May 29 2014;7:40. doi:10.1186/1756-6606-7-40

92. Hamilton A, Vasefi M, Vander Tuin C, McQuaid RJ, Anisman H, Ferguson SS. Chronic Pharmacological mGluR5 Inhibition Prevents Cognitive Impairment and Reduces Pathogenesis in an Alzheimer Disease Mouse Model. Cell Rep. May 31 2016;15(9):1859–65. doi:10.1016/j.celrep.2016.04.077

93. Hu NW, Nicoll AJ, Zhang D, et al. mGlu5 receptors and cellular prion protein mediate amyloid-beta-facilitated synaptic long-term depression in vivo. Nat Commun. Mar 4 2014;5:3374. doi:10.1038/ncomms4374

94. Lee S, Salazar SV, Cox TO, Strittmatter SM. Pyk2 Signaling through Graf1 and RhoA GTPase Is Required for Amyloid-beta Oligomer-Triggered Synapse Loss. J Neurosci. Mar 6 2019;39(10):1910–1929. doi:10.1523/JNEUROSCI.2983-18.2018

95. Zhang D, Qi Y, Klyubin I, et al. Targeting glutamatergic and cellular prion protein mechanisms of amyloid beta-mediated persistent synaptic plasticity disruption: Longitudinal studies. Neuropharmacology. Jul 15 2017;121:231–246. doi:10.1016/j.neuropharm.2017.03.036

96. Kaufman AC, Salazar SV, Haas LT, et al. Fyn inhibition rescues established memory and synapse loss in Alzheimer mice. Ann Neurol. Jun 2015;77(6):953–71. doi:10.1002/ana.24394

97. Lee G, Newman ST, Gard DL, Band H, Panchamoorthy G. Tau interacts with src-family non-receptor tyrosine kinases. J Cell Sci. Nov 1998;111 (Pt 21):3167–77. doi:10.1242/jcs.111.21.3167

98. Ittner LM, Ke YD, Delerue F, et al. Dendritic function of tau mediates amyloid-beta toxicity in Alzheimer’s disease mouse models. Cell. Aug 6 2010;142(3):387–97. doi:10.1016/j.cell.2010.06.036

99. Roberson ED, Halabisky B, Yoo JW, et al. Amyloid-beta/Fyn-induced synaptic, network, and cognitive impairments depend on tau levels in multiple mouse models of Alzheimer’s disease. J Neurosci. Jan 12 2011;31(2):700–11. doi:10.1523/JNEUROSCI.4152-10.2011

100. Larson M, Sherman MA, Amar F, et al. The complex PrP(c)-Fyn couples human oligomeric Abeta with pathological tau changes in Alzheimer’s disease. J Neurosci. Nov 21 2012;32(47):16857–71a. doi:10.1523/JNEUROSCI.1858-12.2012

101. Lambert JC, Ibrahim-Verbaas CA, Harold D, et al. Meta-analysis of 74,046 individuals identifies 11 new susceptibility loci for Alzheimer’s disease. Nat Genet. Dec 2013;45(12):1452–8. doi:10.1038/ng.2802

102. Wang Q, Walsh DM, Rowan MJ, Selkoe DJ, Anwyl R. Block of long-term potentiation by naturally secreted and synthetic amyloid beta-peptide in hippocampal slices is mediated via activation of the kinases c-Jun N-terminal kinase, cyclin-dependent kinase 5, and p38 mitogen-activated protein kinase as well as metabotropic glutamate receptor type 5. J Neurosci. Mar 31 2004;24(13):3370–8. doi:10.1523/JNEUROSCI.1633-03.2004

103. Renner M, Lacor PN, Velasco PT, et al. Deleterious effects of amyloid beta oligomers acting as an extracellular scaffold for mGluR5. Neuron. Jun 10 2010;66(5):739–54. doi:10.1016/j.neuron.2010.04.029

104. Raka F, Di Sebastiano AR, Kulhawy SC, et al. Ca(2+)/calmodulin-dependent protein kinase II interacts with group I metabotropic glutamate and facilitates receptor endocytosis and ERK1/2 signaling: role of beta-amyloid. Mol Brain. Mar 26 2015;8:21. doi:10.1186/s13041-015-0111-4

105. Overk CR, Cartier A, Shaked G, et al. Hippocampal neuronal cells that accumulate alpha-synuclein fragments are more vulnerable to Abeta oligomer toxicity via mGluR5--implications for dementia with Lewy bodies. Mol Neurodegener. May 19 2014;9:18. doi:10.1186/1750-1326-9-18

106. He Y, Wei M, Wu Y, et al. Amyloid beta oligomers suppress excitatory transmitter release via presynaptic depletion of phosphatidylinositol-4,5-bisphosphate. Nat Commun. Mar 13 2019;10(1):1193. doi:10.1038/s41467-019-09114-z

107. Abd-Elrahman KS, Hamilton A, Albaker A, Ferguson SSG. mGluR5 Contribution to Neuropathology in Alzheimer Mice Is Disease Stage-Dependent. ACS Pharmacol Transl Sci. Apr 10 2020;3(2):334–344. doi:10.1021/acsptsci.0c00013

108. Abd-Elrahman KS, Albaker A, de Souza JM, et al. Abeta oligomers induce pathophysiological mGluR5 signaling in Alzheimer’s disease model mice in a sex-selective manner. Sci Signal. Dec 15 2020;13(662) doi:10.1126/scisignal.abd2494

109. Kazim SF, Chuang SC, Zhao W, Wong RK, Bianchi R, Iqbal K. Early-Onset Network Hyperexcitability in Presymptomatic Alzheimer’s Disease Transgenic Mice Is Suppressed by Passive Immunization with Anti-Human APP/Abeta Antibody and by mGluR5 Blockade. Front Aging Neurosci. 2017;9:71. doi:10.3389/fnagi.2017.00071

110. Gregory KJ, Malosh C, Turlington M, et al. Identification of a high affinity MPEP-site silent allosteric modulator (SAM) for the metabotropic glutamate subtype 5 receptor (mGlu(5)). Probe Reports from the NIH Molecular Libraries Program. 2010.

111. Gregory KJ, Dong EN, Meiler J, Conn PJ. Allosteric modulation of metabotropic glutamate receptors: structural insights and therapeutic potential. Neuropharmacology. Jan 2011;60(1):66–81. doi:10.1016/j.neuropharm.2010.07.007

112. Gregory KJ, Noetzel MJ, Rook JM, et al. Investigating metabotropic glutamate receptor 5 allosteric modulator cooperativity, affinity, and agonism: enriching structure-function studies and structure-activity relationships. Mol Pharmacol. Nov 2012;82(5):860–75. doi:10.1124/mol.112.080531

