## Supplementary for "Aging-Dependent Loss of Connectivity in Alzheimer’s Model Mice with Rescue by mGluR5 Modulator"

### Supplementary Material

#### Supplementary Methods

##### Head-plate surgical procedures.

At 3M (N=27), 5/8M (N=4/13), or 19M (N=30), all animals undergo a surgical procedure where an in-house built glass head-plate is affixed to the skull. The surgical procedure is based on the design we have described previously.[34] Since this description, we eliminated the agarose layer and plastic border. We find that these amendments improve the longevity of the preparation. The implants reduce susceptibility artifacts in the cortex and head-motion. The head-plates are well tolerated and the signal-to-noise ratio (SNR) in the cortex shows no change over time (data not shown, non-significant differences in SNR across sessions). The head-plate yields permanent optical access to the cortex (for WF-$Ca^{2+}$ imaging) and immobilizes the head. Animals are initially anesthetized with 5% isoflurane (70/30 medical air/$O_2$) and secured in a stereotaxic frame (KOPF) using ear-bars and an incisor-bar. Isoflurane is then reduced to 2%. Ointment is applied to the eyes to prevent dryness; meloxicam (2mg/kg body weight) is administered subcutaneously, and bupivacaine (0.1%) injected locally at the incision site. After hair removal (Nair), the scalp is washed (3×) with betadine followed by ethanol 70%. The skin and soft tissue overlying the skull surface is surgically removed. The skull is cleaned and dried. Antibiotic powder (Neo-Predef) is applied to the incision site, and isoflurane reduced further to 1.5%. Skull-thinning of the frontal and parietal skull plates is performed with a hand-held drill (FST), tip diameters 1.4 and 0.7mm, to translucency whilst remaining intact. Superglue (Locite) is applied to the exposed skull, followed by transparent dental cement C&B Metabond (Parkell). The pre-cut (Omax 5555 with Tilt-A-Jet, Fisher Finest, Fisher Scientific, PA) head-plate is attached to the dental cement prior to its solidification. Animals are allotted 3-4 weeks to recovered from the procedure before any imaging data are collected.

##### Creation of out-of-sample MRI template.

All operations are executed using BioImage Suite (BIS). We generate the out-of-sample template from N=162 isotropic ($0.2 \times 0.2 \times 0.2 mm^3$) MSME images from N=64 mice (each animal is imaged up to three times during separate imaging sessions with a minimum of 7 days between sessions). Data are skull stripped using an in-house algorithm. After brains are isolated, two images, selected at random, are registered using non-linear registration. The registered images are merged to create an interim reference image. A third image, selected at random from those remaining in the pool, is registered, and merged with the interim reference image using the same approach. This procedure is repeated until no images remain in the pool (i.e., all 162 images are merged). This final merged image is split along the midline and the two halves (hemispheres) are registered and merged. These data are then duplicated, and one 'side' is flipped to create a bilaterally symmetric brain image. Using non-linear registration, this template is registered to the annotated CCFv3 data and ontology downloaded from the Allen Institute.[39,40] We modulated the contrast and smoothed the histological data to better match the MR data prior to this registration step.

#### Structural MRI data acquisition.

Besides the 3D isotropic MSME image, two other structural images are acquired, for registration purposes to the WF-$Ca^{2+}$ data, however these are not used in this study. (2) A multi-spin-multi-echo (MSME) image sharing the same field-of-view (FOV) as the fMRI data. Acquired with a repetition time (TR) of 2500ms and echo time (TE) of 20ms, 28 slices, two averages, and resolution of 0.1×0.1×0.4 $mm^3$ (10mins and 40s). (3) A fast-low-angle-shot (FLASH) time-of-flight (TOF) angiogram (FOV, 2.0×1.0×2.5$cm^3$, covering the cortex), TR/TE of 130/4ms, and resolution of 0.05×0.05×0.05$mm^3$ (18mins). Of note, the FLASH-TOF was not acquired in the animals that did not undergo multimodal imaging (i.e., animals in experiments 2 and 3).

#### Automated skull stripping for template creation.

We use a 2D U-Net-like framework from the deep learning library deep-image. For training data, we manually segmented the brains of five mice selected at random (balanced by session). The binarized masks and MSME images constitute the training set. To make efficient use of the small number of images included in the training data, we implement augmentation methods including image rotation, deformation, cropping, scaling, and padding. Quantile normalization is used for data normalization. Our neural network is based on a three level U-Net architecture. The number of channels in the contracting path are 64, 128 and 256 from top-to-bottom. Each level consists of two padded convolutions with filter size three and stride size one using a ReLU activation function. A max pooling layer of stride size two is applied after the top two levels. In the expansive path, up-convolution of stride size two is conducted between levels. Concatenation is performed by concatenating the up-sampled feature map and the feature map of the corresponding level within the contracting path. Finally, training is performed using the Adam Optimizer with a Cross Entropy loss function.

#### Functional MRI pre-processing and registration to common space

The pre-processed fMRI data from each mouse at each imaging session are averaged to create a representative mean image that is itself corrected for intensity inhomogeneity and then non-linearly registered to the isotropic structural MSME image acquired during the same imaging session. This procedure minimizes the effects of distortions caused by susceptibility artifacts.[75] Then, four transforms -

(1) the framewise rigid head motion correction, (2) representative mean fMRI to individual mouse/session isotropic MSME image, (3) individual mouse/session isotropic MSME image to within-dataset template, and (4) within-dataset template to out-of-sample template – are concatenated and applied to the fMRI data to normalize timeseries in the out-of-sample template space. As part of this step, the native resolution of the functional data ($0.31\times0.31\times0.31mm^3$) is resampled to the template resolution ($0.2\times0.2\times0.2mm^3$). The Allen Atlas (which resides in the out-of-sample template space) can then be applied to the fMRI data to compute connectomes. Registration performance is visually inspected using the RABIES quality control report. Following timeseries normalization, the six-parameters motion estimates are regressed from the timeseries data and used to compute framewise-displacement (FD). Frame scrubbing for motion using a conservative 0.075mm threshold is applied. Cerebrospinal fluid (CSF) and white matter (WM) are also used as nuisance regressors for the final timeseries analyses. Data are smoothed with 0.4mm sigma.

#### Exclusion criteria

From experiment 1 (**Table 1**), N=25 are imaged at the first imaging session (4M) and no animals or data are excluded. Before the second imaging session (6M), N=4 mice (all in the AD group) recovered poorly after the first imaging session and were thus excluded from the 6M imaging session. An additional N=2 were not imaged at the second imaging session (6M) due to scanner hardware issues (but were imaged at the third imaging session). Thus, in total, N=18 are imaged at the second imaging session. Between the second (6M) and third (9M) imaging session, N=9 are allocated to a separate study leaving N=8 for the third imaging session (9M). None of these data are excluded. To augment the number of mice at 6M and 9M (N=18 and N=8, respectively) we conducted experiment 2 (**Table 1**) where N=4 are imaged at 6M and N=13 are imaged at 9M. Thus, between experiment 1 and 2, N=22 at 6M and N=21 at 9M.

For experiment 3, N=27 are imaged at 20M. Of these, N=3 are not scanned at the following imaging session (22M), N=1 due to head-implant detachment, N=1 was euthanized due to an eye infection and N=1 recovered poorly from anaesthesia after the first imaging session. Of note, the animals that experience implant detachment received head-implants (unlike the rest) made with coated glass (Fisher Scientific, same brand as used throughout the other experiments, but with an additional coating that decreased glue-to-glass adherence, thus leading to implant detachment). The remaining N=23 are imaged at 22M, and no animals or data are excluded.

#### Treatment with a silent allosteric mGluR5 (metabotropic glutamate receptor 5) modulator

A subgroup of AD mice (N=10) is treated between 20 and 22M with a silent allosteric mGluR5 (metabotropic glutamate receptor 5) modulator (SAM, BMS-984923). Drug synthesis is as previously described (compound 16)[76] with simplifications that accommodate 5-10 kg scale preparations by Aptuit-Verona [29]. Purity and composition are confirmed to be >97%.[76] The SAM compound is incorporated into OpenStandard Diet with 15% kCal Fat purified diet pellets by Research Diets, Inc. at 50mg SAM per kg food. Vehicle pellets are the same purified diet without any additives (Research Diets, Inc. D11112201i).[29] The drug dosage in food is calculated based on the average amount eaten in a day per kg of body weight[77] and adjusted to be equivalent to ingesting 7.5 mg/kg per day. Previous studies verified plasma levels of BMS-984923 corresponding to those associated with greater than 85% brain mGluR5 receptor occupancy in PET studies. Throughout the treatment period, body weight is monitored to ensure drug/food intake.

#### Triple-network model

From the 172-ROI Allen Atlas, we select ROIs that belong to a DMN based on the description by Whitesell and colleagues.[46] Additionally, ROIs belonging to a SN and LCN are selected based on the triple-network model.[32,33,78] Within our template space, these correspond to: DMN [2:3,18:19,28:31, 38:40], SN [2:3, 28:29, 35:37, 59:61, 63, 70:71, 74:75] and LCN [2:11, 41, 62].

#### Data-derived networks

In this study, we investigate four main contrasts: aging (WT-aging and AD-aging), pathology (AD vs WT) and treatment ($AD_{treated}$ vs $AD_{vehicle}$). For each of these, we generate a 'data-derived' network: a binarized mask of the 172x172 matrix, where each value=1/-1 corresponds to all surviving positive/negative edges from the t-tests (positive/negative t-values, respectively generating a positive network and a negative network) computed for that specific condition (see **Fig. 2, 4 and 5** and **Statistics section**). For example, the WT-aging network is obtained by (first) computing t-tests at 4M>9M and 4M>20M timepoints, and then creating binarized matrices for both t-test results. Finally, the two matrices obtained (two positive and two negative) are combined either by sum or by intersect to obtain a more extensive or a more stringent data-derived network.

### Supplementary Discussion

#### BMS-984923 mechanism of action

The cellular prion protein (PrP$^C$) is a specific and high affinity receptor for amyloid-β oligomers (Aβo).[79-86] mGluR5 links the Aβo/PrP$^C$ complex to the intracellular tyrosine kinases Fyn (a neuronally-enriched cytoplasmic tyrosine kinase) and Pyk2 (PTK2B).[31,85,87-95] These kinases are in turn associated with tau,[96-100] and are implicated in AD-risk.[101] At the nexus of this AD-pathway, mGluR5 is a promising target. Indeed, reduction of mGluR5 activity alleviates synaptic and memory deficits in multiple AD models.[48,87,89-93,102-109] Further, BMS-984923 spares glutamate signaling[110-112] while still inhibiting the mGluR5 activation by pathological Aβo/PrP$^C$ complexes.[31] This mechanism of action greatly expands the potential therapeutic window for mGluR5 as a disease-modifying target.

### Supplementary Figures

A.

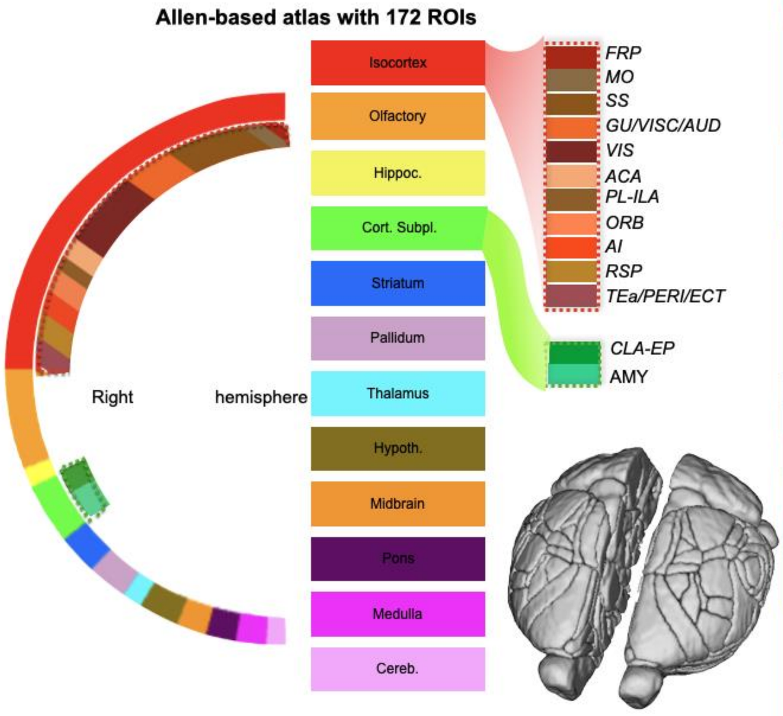


B.

**Atlas and RSNs breakdown**

| | | |
|---|---|---|
| R/L-FRP | 1: R/L-Frontal pole, cerebral cortex | |
| R/L-MOp | 2: R/L-Primary motor area | DMN/SN/LCN |
| R/L-MOs | 3: R/L-Secondary motor area | |
| R/L-SSp-n | 4: R/L-Primary somatosensory area, nose | LCN |
| R/L-SSp-bfd | 5: R/L-Primary somatosensory area, barrel field | |
| R/L-SSp-ll | 6: R/L-Primary somatosensory area, lower limb | |
| R/L-SSp-m | 7: R/L-Primary somatosensory area, mouth | |
| R/L-SSp-ul | 8: R/L-Primary somatosensory area, upper limb | |
| R/L-SSp-tr | 9: R/L-Primary somatosensory area, trunk | |
| R/L-SSp-un | 10: R/L-Primary somatosensory area, unassigned | |
| R/L-SSs | 11: R/L-Supplemental somatosensory area | |
| R/L-GU | 12: R/L-Gustatory areas | |
| R/L-VISC | 13: R/L-Visceral area | |
| R/L-AUDd | 14: R/L-Dorsal auditory area | |
| R/L-AUDp | 15: R/L-Primary auditory area | |
| R/L-AUDpo | 16: R/L-Posterior auditory area | |
| R/L-AUDv | 17: R/L-Ventral auditory area | |
| R/L-VISal | 18: R/L-Anterolateral visual area | DMN |
| R/L-VISam | 19: R/L-Anteromedial visual area | |
| R/L-VISl | 20: R/L-Lateral visual area | |
| R/L-VISp | 21: R/L-Primary visual area | |
| R/L-VISpl | 22: R/L-Posterolateral visual area | |
| R/L-VISpm | 23: R/L-posteromedial visual area | |
| R/L-VISa | 24: R/L-Anterior area | |
| R/L-VISli | 25: R/L-Laterointermediate area | |
| R/L-VISrl | 26: R/L-Rostrolateral visual area | |
| R/L-VISpor | 27: R/L-Postrhinal area | |
| R/L-ACAd | 28: R/L-Anterior cingulate area, dorsal part | DMN/SN |
| R/L-ACAv | 29: R/L-Anterior cingulate area, ventral part | |
| R/L-PL | 30: R/L-Prelimbic area | DMN |
| R/L-ILA | 31: R/L-Infralimbic area | |
| R/L-ORBl | 32: R/L-Orbital area, lateral part | |
| R/L-ORBm | 33: R/L-Orbital area, medial part | |
| R/L-ORBvl | 34: R/L-Orbital area, ventrolateral part | |
| R/L-AId | 35: R/L-Agranular insular area, dorsal part | SN |
| R/L-AIp | 36: R/L-Agranular insular area, posterior part | |
| R/L-AIv | 37: R/L-Agranular insular area, ventral part | |
| R/L-RSPagl | 38: R/L-Retrosplenial area, lateral agranular part | DMN |
| R/L-RSPd | 39: R/L-Retrosplenial area, dorsal part | |
| R/L-RSPv | 40: R/L-Retrosplenial area, ventral part | |
| R/L-TEa | 41: R/L-Temporal association areas | LCN |
| R/L-PERI | 42: R/L-Perirhinal area | |
| R/L-ECT | 43: R/L-Ectorhinal area | |
| R/L-MOB | 44: R/L-Main olfactory bulb | |
| R/L-AOB | 45: R/L-Accessory olfactory bulb | |
| R/L-AON | 46: R/L-Anterior olfactory nucleus | |
| R/L-TT | 47: R/L-Taenia tecta | |
| R/L-DP | 48: R/L-Dorsal peduncular area | |
| R/L-PIR | 49: R/L-Piriform area | |
| R/L-NLOT | 50: R/L-Nucleus of the lateral olfactory tract | |
| R/L-COA | 51: R/L-Cortical amygdalar area | |
| R/L-PAA | 52: R/L-Piriform-amygdalar area | |
| R/L-TR | 53: R/L-Postpiriform transition area | |
| R/L-HIP | 54: R/L-Hippocampal region | |
| R/L-RHP | 55: R/L-Retrohippocampal region | |
| R/L-CLA | 56: R/L-Claustrum | |
| R/L-EP | 57: R/L-Endopiriform nucleus | |
| R/L-LA | 58: R/L-Lateral amygdalar nucleus | |
| R/L-BLA | 59: R/L-Basolateral amygdalar nucleus | SN |
| R/L-BMA | 60: R/L-Basomedial amygdalar nucleus | |
| R/L-PA | 61: R/L-Posterior amygdalar nucleus | |
| R/L-STRd | 62: R/L-Striatum dorsal region | LCN |
| R/L-STRv | 63: R/L-Striatum ventral region | SN |
| R/L-LSX | 64: R/L-Lateral septal complex | |
| R/L-sAMY | 65: R/L-Striatum-like amygdalar nuclei | |
| R/L-PALd | 66: R/L-Pallidum, dorsal region | |
| R/L-PALv | 67: R/L-Pallidum, ventral region | |
| R/L-PALm | 68: R/L-Pallidum, medial region | |
| R/L-PALc | 69: R/L-Pallidum, caudal region | |
| R/L-DORsm | 70: R/L-Thalamus, sensory-motor cortex related | SN |
| R/L-DORpm | 71: R/L-Thalamus, polymodal association cortex related | |
| R/L-PVZ | 72: R/L-Periventricular zone | |
| R/L-PVR | 73: R/L-Periventricular region | |
| R/L-MEZ | 74: R/L-Hypothalamic medial zone | SN |
| R/L-LZ | 75: R/L-Hypothalamic lateral zone | |
| R/L-MBsen | 76: R/L-Midbrain, sensory related | |
| R/L-MBmot | 77: R/L-Midbrain, motor related | |
| R/L-MBsta | 78: R/L-Midbrain, behavioral state related | |
| R/L-P-sen | 79: R/L-Pons, sensory related | |
| R/L-P-mot | 80: R/L-Pons, motor related | |
| R/L-P-sat | 81: R/L-Pons, behavioral state related | |
| R/L-MY-sen | 82: R/L-Medulla, sensory related | |
| R/L-MY-mot | 83: R/L-Medulla, motor related | |
| R/L-MY-sat | 84: R/L-Medulla, behavioral state related | |
| R/L-CBX | 85: R/L-Cerebellar cortex | |
| R/L-CBN | 86: R/L-Cerebellar nuclei | |

**Fig. S1| 172-ROIs from the Allen Atlas and their breakdown into the triple-network system.** (**A.**) Breakdown of (rainbow) color-coding of brain regions used in this work (main figures). (**B.**) Allen Atlas identities of the 172 regions we use in this work and their membership within the triple network system: default-mode network, salience network and latero-cortical network (DMN, SN, LCN). Isocortex = FRP: frontal pole, MO: motor areas, SS: somatosensory areas, GU: gustatory area, VISC: visceral area, AUD: auditory area, VIS: visual area, ACA: anterior cingulate area, PL: prelimbic area, ILA: infralimbic area, ORB: orbital area, AI: agranular insular area, RSP: retrosplenial areas, TEa: temporal association areas, PERI: perirhinal area, ECT: ectorhinal area, OLF: olfactory areas, HF: hippocampal formation; Cortical Subplate = CLA: claustrum, EP: Endopiriform nucleus, AMY: amygdala; STR: striatum, PAL: pallidum, TH: thalamus, HT: hypothalamus, MB: midbrain, P: pons, MED: medulla, CBX: cerebellar cortex.

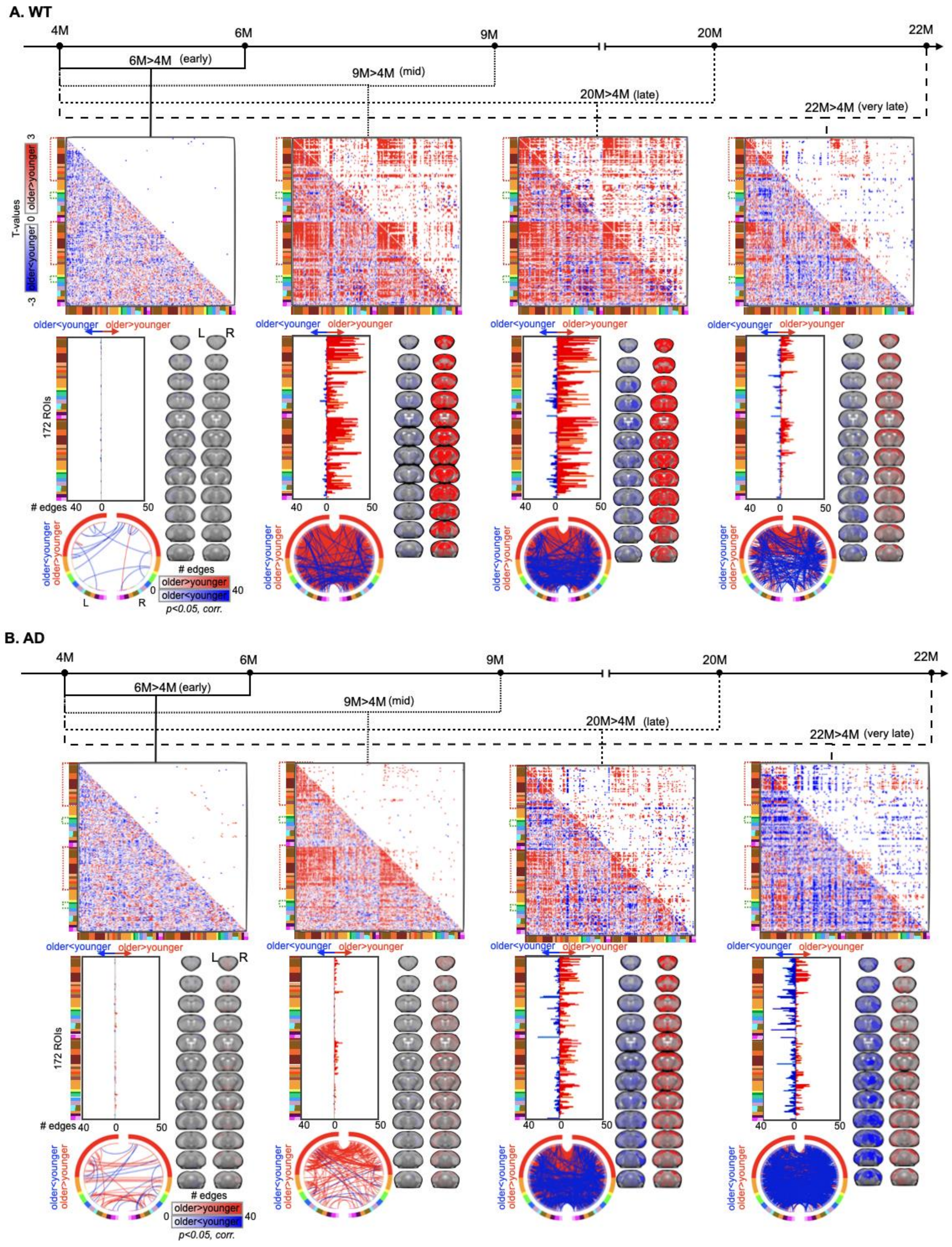


***Caption in the next page***

**Fig. S2| WT and AD follow distinct trajectories of age-related changes in functional brain organization.** As in **Fig. 2** except here, we use all data (from WT and AD mice) as the 4M baseline for both aging trajectories. The results for both WT (**A.**) and AD (**B.**) are similar to those obtained within either group (see main text, **Fig. 2**).

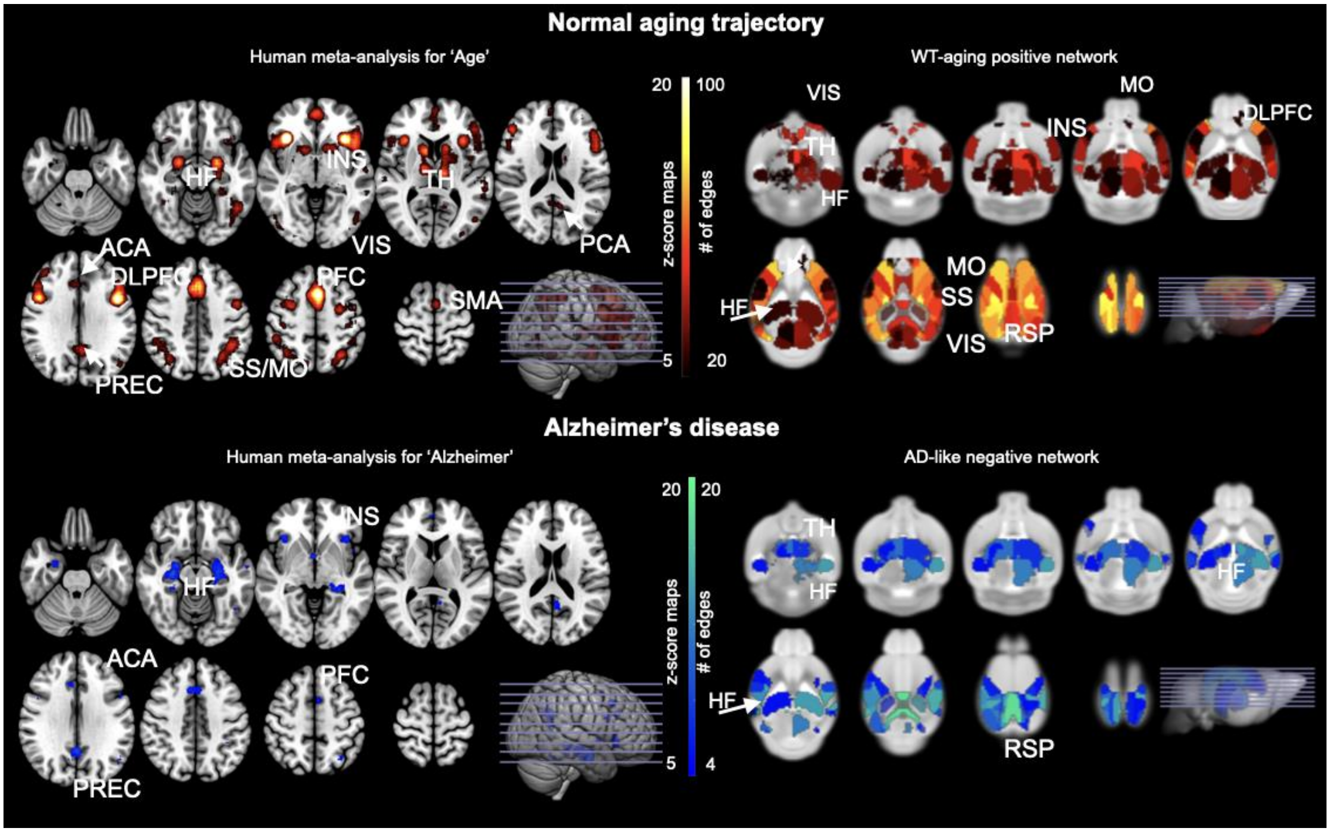


**Fig. S3| Comparison of WT-aging and AD-like networks with human meta-analyses.** We have utilized the online platform for automated synthesis of fMRI studies Neurosynth (http://neurosynth.org) to compare the spatial structure of our AD-network to human neuroimaging studies using the queries 'Age' and 'Alzheimer'. From the Neurosynth database we have downloaded the association test map (z-score maps, corrected for false discovery rate (FDR = 0.01)),[47] for both Age-map (top, left) and Alzheimer (bottom, left), in red and blue hues, respectively. Here, we compare these to the WT-aging (positive, top-right) and AD-like (negative, bottom-right) network profiles in the mouse. Although the meta-analysis reported does not provide information on the directionality of the activation in either case, there is an interesting overlap in the spatial distribution of the maps for both aging and Alzheimer, with clear overlap with our mouse-derived networks. Specifically, 'Age' query identifies cortical regions to be highly involved, and regions like the cingulate cortex, precuneus and prefrontal areas; these highly overlap with the WT-aging network in the mouse. The query for 'Alzheimer' shows higher involvement of AD-vulnerable brain regions such as the HF, which is also reported in AD-like pathology. HF: hippocampal formation; PREC: precuneus; ACA: anterior cingulate areas; INS: insula; TH: thalamus; PFC: prefrontal cortex; RSP: retrosplenial cortex; VIS: visual areas; SMA: supplementary motor area; PCA: posterior caudate; SS: somatosensory; MO: motor; DLPFC: dorsolateral prefrontal cortex.

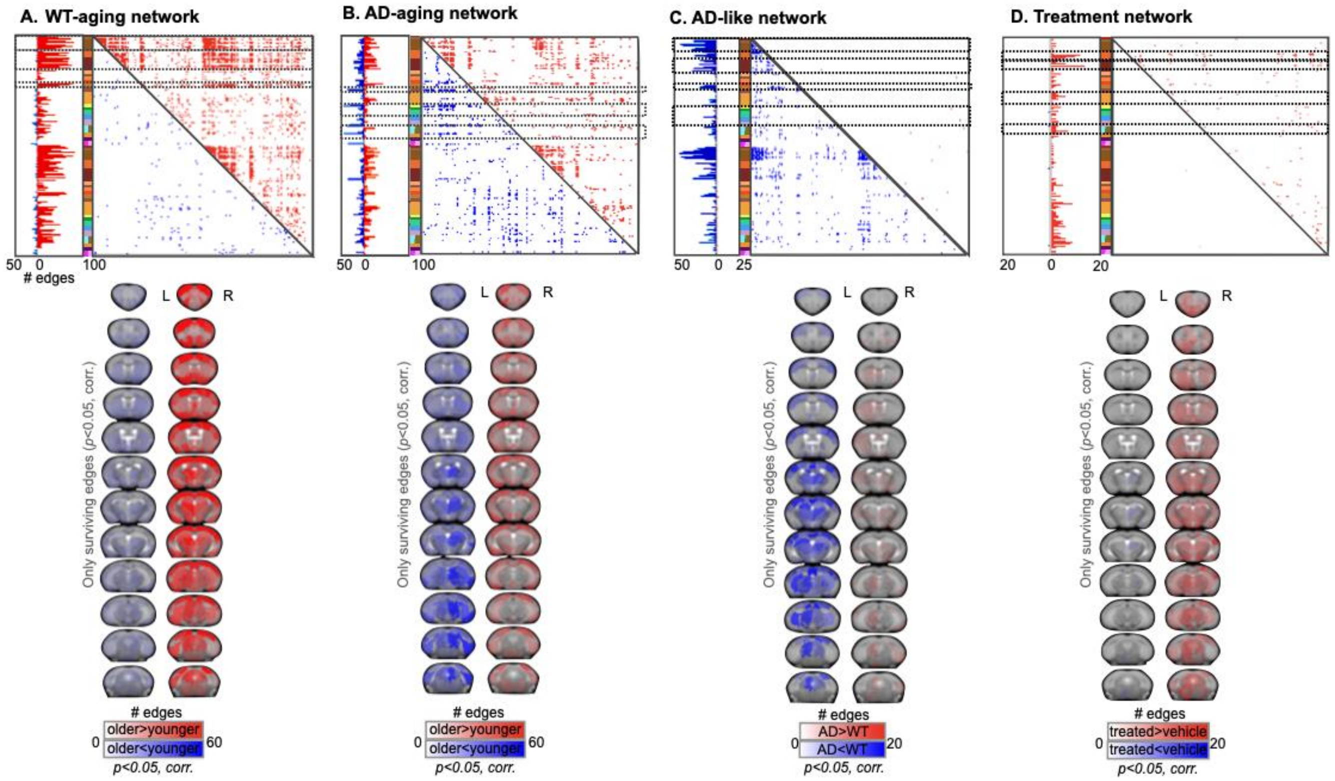


**Fig. S4| Summary of data-derived networks.** Summary reproduction of each of the four data-derived networks: WT and AD-aging (**A., B.,** respectively), and AD-like and treatment networks (**C., D.,** respectively). Here the sum (less stringent) is adopted.

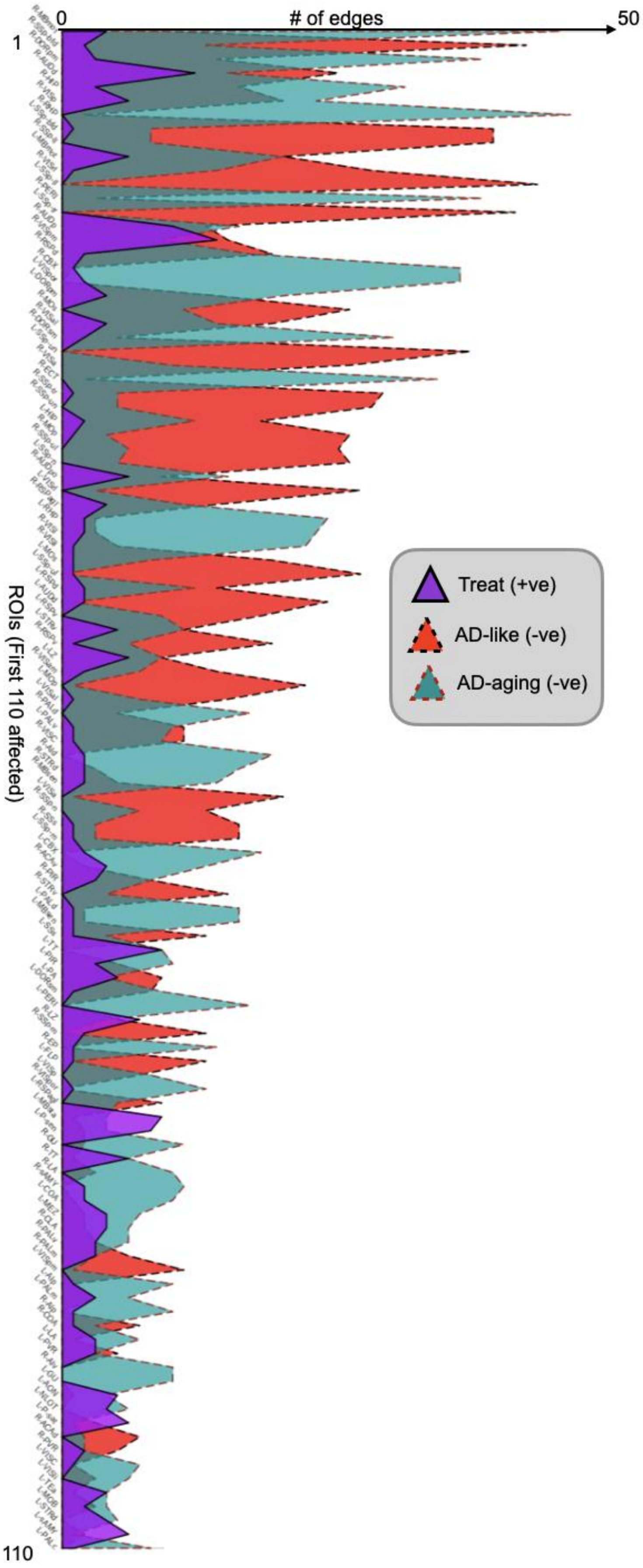


**Fig. S5| Summary of data-derived networks.** To assess the specificity of the treatment in the regions affected both in AD-aging and AD-like pathology, the node-degree maps for AD-aging and AD-like negative edges (-ve) and treatment positive (+ve) are compared in a layered area graph, thresholding to the first 110 regions involved.

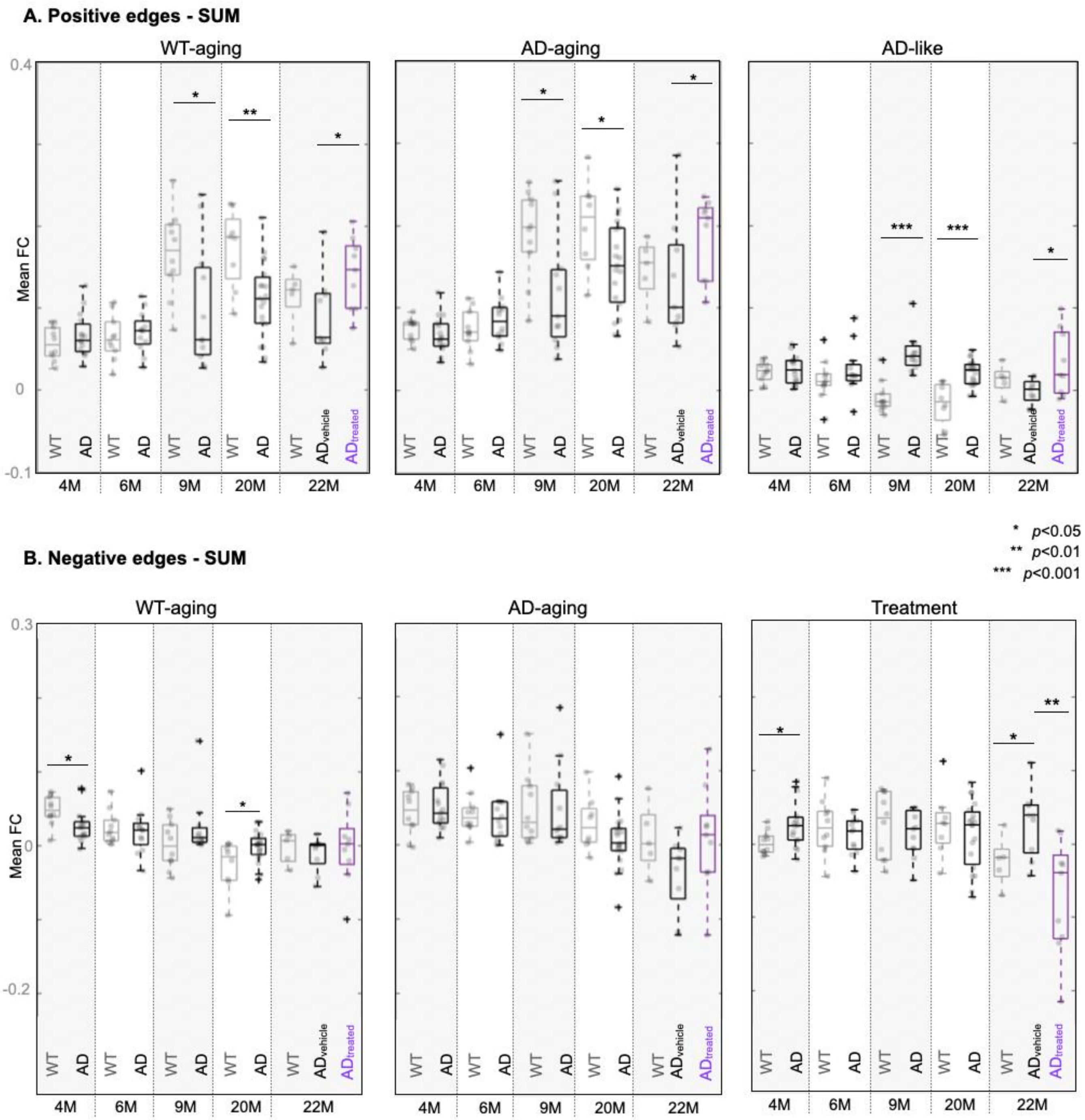


**Fig. S6| Mean FC within data-derived networks (sum).** Of the four data-derived networks (reproduced in **Fig. S4**, for the sum), the positive networks for WT-aging, AD-aging and AD-like (**A.**), and the negative networks for WT-aging, AD-aging and treatment (**B.**) are used as binary masks to threshold the raw functional connectome data, and to plot FC values corresponding to each network. As such, the functional connectomes are thresholded for only the edges belonging to each of the selected networks, for all time points and for both WT (grey), AD (vehicle/treated black/dark red) animals, for negative (**A.**) and positive (**B.**) edges. The correlation values within each run are summed, and the three runs per mouse averaged. Each data point in the boxplots above represent a mouse average. The mean FC values are used in a t-test to compare across genotypes, at each time point. Horizontal lines show the median within each group. Significant differences between WT and AD mice are reported as * for $p<0.05$, ** for $p<0.01$, and *** for $p<0.001$. Mean FC values are shown for positive and negative-edges networks (**A., B.,** respectively).
